# Concerted transformation of a hyper-paused transcription complex and its reinforcing protein

**DOI:** 10.1101/2023.06.26.546588

**Authors:** Philipp K. Zuber, Nelly Said, Tarek Hilal, Bing Wang, Bernhard Loll, Jorge González-Higueras, César A. Ramírez-Sarmiento, Georgiy A. Belogurov, Irina Artsimovitch, Markus C. Wahl, Stefan Knauer

## Abstract

RfaH, a paralog of the universally conserved NusG, binds to RNA polymerases (RNAP) and ribosomes to activate expression of virulence genes. In free, autoinhibited RfaH, an α-helical KOW domain sequesters the RNAP-binding site. Upon recruitment to RNAP paused at an *ops* site, KOW is released and refolds into a β-barrel, which binds the ribosome. Our structures of *ops*-paused transcription elongation complexes alone and bound to the autoinhibited and activated RfaH reveal swiveled, pre-translocated pause states stabilized by an *ops* hairpin in the non-template DNA. Autoinhibited RfaH binds and twists the *ops* hairpin, expanding the RNA:DNA hybrid to 11 base pairs and triggering the KOW release. Once activated, RfaH hyper-stabilizes the pause, which thus requires anti-backtracking factors for escape. Our results suggest that the entire RfaH cycle is solely determined by the *ops* and RfaH sequences and provide insights into mechanisms of recruitment and metamorphosis of NusG homologs across all life.

**HIGHLIGHTS:**

- The nontemplate DNA strand of an *ops*-paused transcription complex forms a hairpin
- Autoinhibited RfaH binds and twists the *ops* hairpin to expand the RNA:DNA hybrid
- RfaH-hairpin contacts are solely responsible for triggering RfaH activation
- Upon recruitment, RfaH hyper-stabilizes the pause and promotes RNAP backtracking

## INTRODUCTION

In every cell, RNA synthesis is modulated by a wide array of accessory proteins that bind to multi-subunit RNA polymerase (RNAP) and nucleic acids and adjust gene expression to cellular demands. Among these transcription factors, NusG (Spt5/DSIF in archaea and eukaryotes) stands out as the only regulator conserved across all life ^1^. NusG proteins bind to RNAP genome-wide ^2, 3^ to promote efficient synthesis and folding of the nascent RNA ^4–6^ and consist of a NusG N-terminal (NGN) domain flexibly connected to one (or several in eukaryotes) C-terminal Kyprides, Ouzounis, Woese (KOW) domains. The NGNs share α/β topology, bind to a conserved site on the largest subunit of RNAP, and are sufficient for all direct effects on RNA synthesis ^5, 7–12^. The β-barrel KOW domains contact diverse proteins to couple transcription to RNA folding, modification, splicing, nucleosome remodeling, translation, and other cellular processes ^6, 13–20^.

Many cellular genomes encode specialized paralogs of NusG ^21^, whose roles are best characterized in bacteria, where they are required for biosynthesis of capsules ^22, 23^, toxins and adhesins ^24, 25^, antibiotics ^26, 27^, and lipopolysaccharides ^28, 29^. Many NusG paralogs are encoded on conjugative plasmids, from F-factor ^30^ to multidrug-resistant clinical isolates ^31^. Thus, NusG paralogs are vital for bacterial fitness and pathogenesis and may also facilitate evolution ^32^.

NusG paralogs function alongside NusG and have just a few targets, which become critical in some conditions, *e.g.,* during *Klebsiella pneumoniae* lung infection ^22^. The regulatory logic that underpins the division of labor among NusG-like proteins is well understood in *Escherichia coli.* NusG is an abundant protein that dynamically interacts with almost every transcribing RNAP ^3^ and determines the fate of the nascent RNA by either suppressing or promoting Rho-dependent termination. On translated mRNAs, NusG can bridge RNAP to the leading ribosome ^19, 20^ whereas on rRNA, NusG is part of an antitermination complex ^6^; in both cases, the nascent RNA and RNAP are shielded from Rho. On antisense, aberrant, and xenogeneic RNAs, NusG KOW binds to Rho to induce premature termination ^33–35^. Xenogeneic silencing is an essential function of *E. coli* NusG ^36^.

Conversely, expression of several virulence and housekeeping operons critically depends on NusG paralog RfaH ^37^. RfaH, but not NusG, is associated with RNAP transcribing these xenogeneic operons ^38^, even though NusG vastly outnumbers RfaH in the cell ^39^. A combination of sequence-specific recruitment and fold-switching-controlled autoinhibition ensures that RfaH finds its targets while not compromising the essential function of NusG ^32^. In free RfaH, the RNAP-binding site on NGN is masked by its KOW domain adopting an α-helical hairpin (KOW^α^) ^11^. RfaH recruitment requires a 12-nucleotide (nt) operon polarity suppressor (*ops*) sequence (Figure 1) present in 5’-untranslated regions of RfaH target operons ^28^. The *ops* element halts RNAP to allow time for RfaH recruitment ^40^ and makes direct contacts to RfaH NGN ^12, 41^. Upon binding to the *ops-*paused elongation complex (*ops*PEC), RfaH is activated through domain dissociation ^42^. The released NGN accommodates on RNAP and converts the enzyme into a pause-resistant state ^12^, while the freed KOW refolds into a NusG-type five-stranded β-barrel (KOW^β^) and binds to the ribosomal protein S10 ^42^ to couple transcription to translation and activate protein synthesis ^43^. Notably, RfaH recruitment to RNAP and the ribosome must be tightly orchestrated: *ops* is the only chance for RfaH to load onto transcribing RNAP, and the ribosome must be captured by RfaH between *ops* and the start codon, located within 100 nts downstream. Failure of either recruitment cripples gene expression by hundreds of folds ^37, 43^.

**Figure 1.**
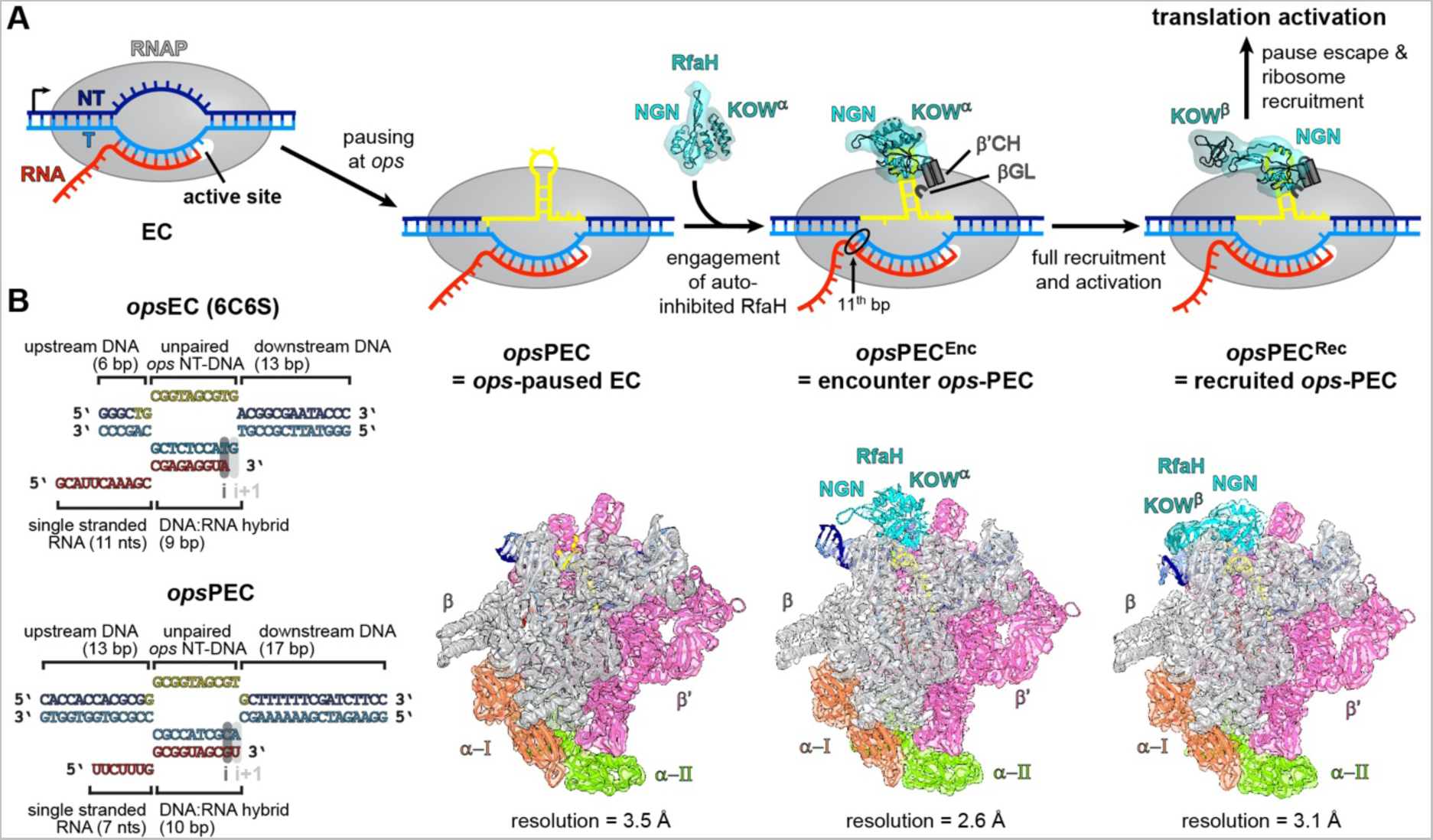
Steps in RfaH recruitment and activation. **(A)** Mechanisms of RNAP pausing at *ops* and RfaH recruitment revealed by cryoEM structures. In *ops*PEC, the NT-DNA strand folds into a hairpin that recruits autoinhibited RfaH to form the PEC^Enc^. During subsequent RfaH activation, KOW^α^ is released from NGN, which establishes stable contacts with RNAP in the resulting *ops*PEC^Rec^, while KOW^α^ refolds into KOW^β^ to set up a stage for recruitment of a ribosome to initiate translation. The cryoEM densities (transparent surfaces) and accompanying models (cartoons) of *ops*PEC, *ops*PEC^Enc^, and *ops*PEC^Rec^ are shown below. (B) Nucleic acids scaffolds used for the assembly of a post-translocated *ops*EC ^12^ (top) and the pre-translocated *ops*PEC used in this study (bottom). In this and other figures, the NT-DNA is shown in dark blue, the T-DNA in light blue, the *ops* element in yellow, and RNA in red.

Understanding such synchronicity requires elucidation of minute structural detail. While much insight has been provided by existing structures of autoinhibited and activated RfaH (in isolation, bound to *ops* DNA, and bound to an *ops*-modified non-paused EC ^11, 12, 41, 42^), their limitations leave several key questions unanswered. First, what features of *ops*PEC render it exceptionally efficient at recruiting RfaH, which is present at fewer than 100 copies per cell? Second, in the autoinhibited RfaH, the RNAP-binding site is partially occluded – how does RfaH bind to *ops*PEC? Third, how is RfaH domain dissociation triggered upon binding to *ops*PEC? Fourth, after accommodation of RfaH, how are ribosome recruitment and pause escape achieved? In this work, we sought to answer these questions by comparing structures of *ops*PECs alone and bound to the autoinhibited and activated RfaH. We show that the formation of an *ops* hairpin (*ops*HP) in the non-template (NT) DNA strand stabilizes a swiveled, pre-translocated paused state with an extended bubble and a 10-base pair (bp) RNA:DNA hybrid. Using the *ops*HP as a handle, autoinhibited RfaH docks onto *ops*PEC near its final binding site, priming the KOW for activation. RfaH grabs and twists the *ops*HP, further expanding the bubble and the hybrid to 11 base pair (bp) and thus hyper-stabilizing a pause that requires accessory factors for escape. Finally, our molecular dynamics simulations suggest that, following the initial domain separation, the RNAP-RfaH contacts are dispensable for the KOW α-to-β fold-switch. Together, our findings illuminate the molecular details underlying a paradigmatic, multi-step process of deoxyriboregulation of RNAP.

## RESULTS

### RNAP pauses at the *ops* element in a pre-translocated state

To enable RfaH recruitment, RNAP (a five-subunit α_2_ββ’ω complex in *E. coli*) pauses at the *ops* site, yielding a pre-translocated PEC ^44^. A previous analysis revealed the structure of a non-paused EC bound to RfaH (RfaH-*ops*EC) ^12^. In that study, the EC was assembled on a scaffold (Figures 1B and S2A) with the *ops* element in the NT DNA, which is sufficient for RfaH binding ^40^, and a template (T) DNA strand that lacked complementarity in the last 10 *ops* nucleotides, most notably the 3’-terminal G12 in the NT DNA (G12; thereafter, all nucleotides are numbered to reflect their positions in the *ops* element, with RNA and T-DNA nts denoted with an R/T superscript), thereby favoring the separation of G12 from the T DNA and the adoption of a post-translocated state. Furthermore, the RNA sequence allowed only a 9-bp hybrid, whereas longer hybrids may occur during *ops*-mediated and RfaH-stabilized pausing. Finally, the scaffold had a short 6-bp upstream DNA duplex, possibly precluding nucleic acid repositioning *via* RNAP or factor contacts to more distal DNA regions. Thus, although the structure of RfaH-*ops*EC captured the molecular details of RfaH interactions with RNAP and DNA, the molecular basis of initial *ops*-mediated pausing is presently unknown.

Here, we assembled an EC on a fully complementary scaffold harboring the *ops* site, an extended upstream duplex (up to 14 bps), and an RNA that could form a hybrid of up to 11 bps at the expense of the proximal upstream DNA melting (Figure 1B). We then elucidated the atomic structure of thus assembled *ops*PEC by cryogenic electron microscopy (cryoEM) in combination with single-particle analysis (SPA) (Figures 1 and S1, Table S1). The particles were highly homogeneous and in the cryoEM reconstruction, almost all regions of RNAP, all DNA nts and 10 nts of RNA were well resolved, with fragmented density for RNA outside of the RNA exit channel.

A PEC formed at *ops* is biochemically distinct from a PEC stabilized by an RNA hairpin, e.g., *his*PEC ^44^. Consistently, while in structures of *his*PECs, the RNA:DNA hybrid adopted a half-translocated state ^12, 45^ (RNA post-translocated, DNA pre-translocated), our *ops*PEC resides in the pre-translocated state (Figure 2A). RNAP is paused at U11, and no unpaired T-DNA nucleotide is present in the i+1 position to receive an incoming rNTP. Compared to an elongation-competent, post-translocated EC ^46^, *his*PECs adopt a swiveled state, in which a so-called swivel module (clamp, dock, shelf, SI3, and a C-terminal segment of the β’ subunit) is rotated by ∼3 ° about an axis perpendicular to the plane defined by the axes of the upstream DNA duplex and the RNA:DNA hybrid ^12, 45^. Swiveling is thought to stabilize the paused state by counteracting folding of the β’ trigger loop (TL), which is required for nucleotide addition ^12, 45^. Consistent with the exceptionally strong pausing at *ops* ^47^, *ops*PEC undergoes particularly pronounced swiveling of 5.8° (Figure 2B). As expected for a swiveled state, the TL is unfolded (Figure 2C) and the β’ SI3 domain is in the open conformation. In *ops*PEC, but not in EC ^46^, the TL β’R933 forms salt bridges with βE546 and βD549, an interaction that may stabilize the unfolded TL (Figure 2C).

**Figure 2.**
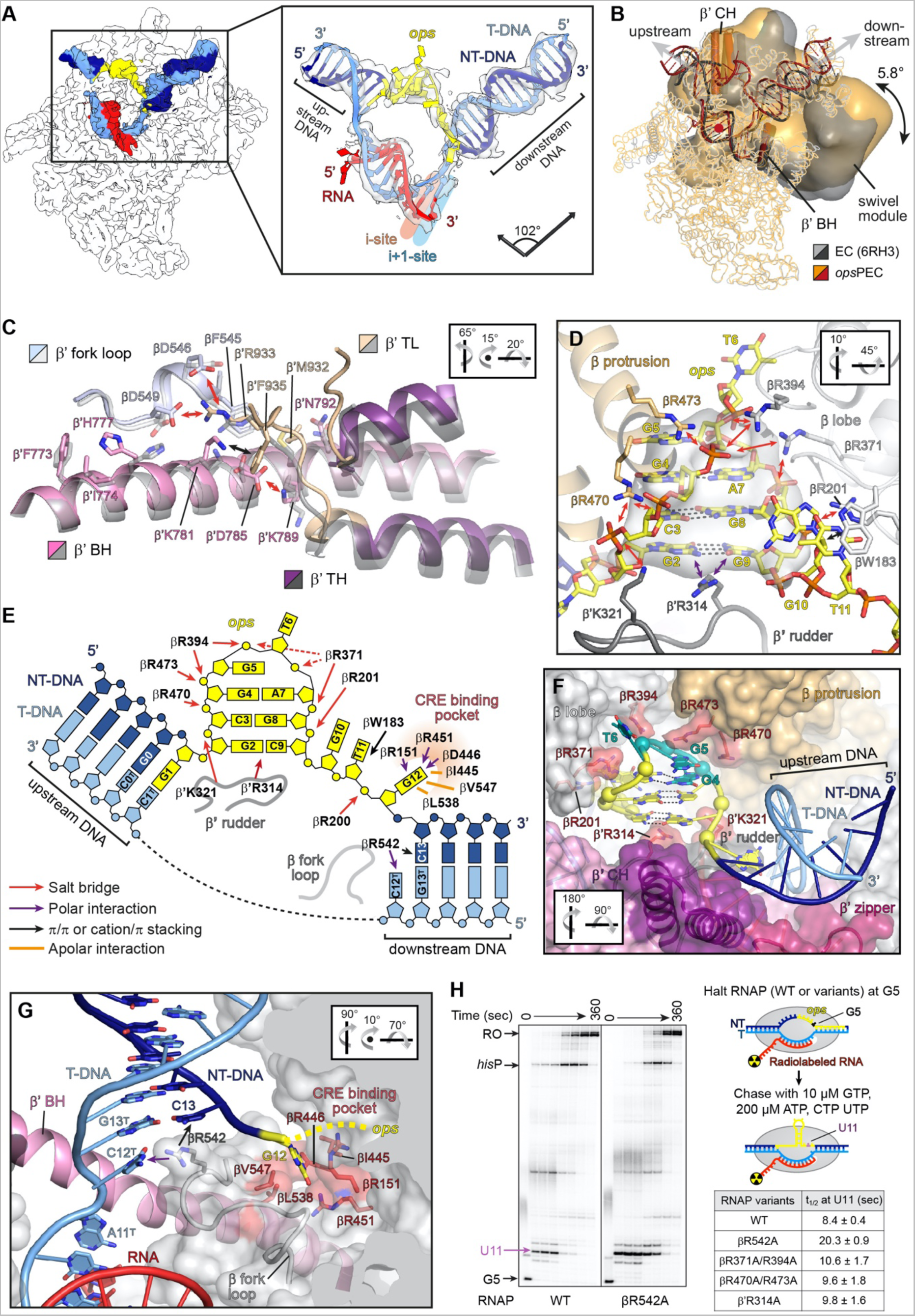
Structural basis of transcriptional pausing in the *ops*PEC. (A) Left: CryoEM map of RNAP is shown as a white outline, nucleic acids maps – as solid surface. Right: Model of the nucleic acid scaffold (cartoon) with its associated cryoEM density (transparent surface). In this and other figures, the arrows represent the helix vectors of the up- and downstream DNA duplexes, and the angle between them is given. (B) A structural overlay of a pre-translocated EC (PDB ID 6RH3) and *ops*PEC. superimposed on the core module (mainly α- and β-subunits; shown as ribbons); the swivel module is represented as Gaussian surface. The swivelling angle and the approximate rotation axis (red dot) is shown. The β’BH and β’CH are displayed as cartoon tubes to illustrate the orientation of the swivelling axis (parallel to β’BH). (C) Superposition of β’BH and β’TL of the *ops*PEC and a post-translocated EC (PDB ID 6ALF; grey). Arrows indicate interactions within residues (sticks) that trap the inactive β’TL. (D) The *ops* DNA hairpin in the NT strand. The *ops*HP and side chains of interacting residues are shown as sticks, the RNAP elements – as transparent cartoons. The cryoEM density is shown as grey surface, H-bonds formed by the *ops* bps – as dashed lines. (E) The RNAP:*ops* interactions. DNA nucleotides are depicted as blocks (bases), rings (ribose), or circles (phosphates), respectively. Salt bridges, polar or apolar interactions with selected RNAP residues are color-coded and indicated by arrows or lines. (F) Stabilization and accessibility of the *ops*HP within the main channel cleft. RNAP is shown as transparent surface; selected structural elements – as coloured cartoons. Side chains of *ops*-interacting residues are in salmon, the *ops* nts interacting with RfaH – in mint. (G) Separation of the *ops* G12:C12^T^ bp at the downstream DNA fork junction. RNAP (sliced at the active site; β’ is omitted for clarity) is shown as transparent surface, nucleic acids as cartoon/sticks. G12 is inserted into the CRE pocket (red), whereas C12^T^ is stabilized by β fork loop 2; red arrows indicate interactions with βR542. In panels C, D, F, and G, orientations relative to panel A are indicated. (A) (H) Effects of RNAP residue substitutions on pausing. Halted radiolabeled G5 ECs were formed with wt or mutationally-altered RNAPs. Single-round elongation assays were carried out as described in STAR Methods. Samples withdrawn at 0, 10, 20, 30, 60, 90, 180, and 360 sec were analyzed on a urea-acrylamide gel; a representative gel is shown. The positions of *ops* G5 and U11, *his*P, and run-off (RO) RNAs are indicated with arrows. The half-life (t_1/2_) of pausing at U11 is presented as mean ± SD; n=3.

In contrast to all other structures of factor-free ECs reported so far, the NT strand is fully defined in the cryoEM reconstruction (Figure 2A). NT bases G2-C9 form a hairpin that rests on top of the R314 side chain of the β’ rudder (Figures 2D and 2E). The stem of the *ops*HP comprises two canonical Watson-Crick (WC) bps (G2:C9 and C3:G8) and a Saenger XI bp (G4:A7) and is stabilized by positively charged residues of the β lobe (R201, R371, R394), β protrusion (R470, R473), and β’ rudder (K321); G1 forms the most proximal bp of the upstream DNA duplex (Figure 2E). Accommodation of the *ops*HP at the β lobe/protrusion pushes the upstream DNA duplex away from the β protrusion and against the β’ zipper and the β’ clamp helices (CH), effectively promoting swiveling (Figures 2B and 2F). Consequently, the upstream and downstream DNA duplexes span an angle of 102 °, vs ∼129 ° in the canonical EC (Figures 2A and S2B).

The two-nt *ops*HP loop (G5-T6) is required for specific recognition by RfaH ^12, 41^. In *ops*PEC, this loop is located between the upstream DNA channel and the main channel of RNAP (Figure 2F). The T6 base is completely flipped outwards and is highly flexible, as indicated by the lack of cryoEM density (Figure 2A), and G5 is also rotated outwards. The *ops*HP-induced displacement of the upstream DNA and swiveling lead to an increased solvent accessibility of the *ops*HP loop, presenting the nucleobases of G5 and T6 for sequence-specific readout.

Immediately downstream of the *ops*HP, the NT DNA changes direction; G10 and T11 move away from the transcription bubble and form an extended stack with βW183 that is laterally stabilized by βD199 and βR200 (β lobe; Figures 2D and 2E). G12 is embedded in the core recognition element (CRE) pocket ^48^, between βR151, βI445, βD446, βR451, βL538, and βV547 (β lobe, β protrusion, and following residues; Figure 2G). Thus, despite the presence of a complementary C12^T^, sequestration of G12 at the CRE prevents it from pairing with C12^T^; instead, the following C13 is diverted to form the first bp of the downstream DNA duplex (Figure 2G).

In the canonical pre-translocated EC, 10 DNA bps are melted to form a 10-bp RNA:DNA hybrid ^49^. Formation of the *ops*HP requires 11 single-stranded (ss) nts in the NT strand – thus, an additional upstream DNA bp is melted and the last unpaired T-strand nt, C12^T^, remains stacked on the downstream duplex and does not move into the templating i+1 position (Figure 2G). Consequently, the DNA strands in the bubble are compressed as compared to canonical ECs, causing a sharper angle between the upstream and downstream duplexes.

The side chain of βR542 occupies the position of G12 in the duplex, engaging the Watson-Crick face of C12^T^ (Figure 2G). This interaction may stabilize the pre-translocated state or promote local melting of downstream DNA to facilitate pause escape. To decipher the βR542 contribution, we substituted it for an alanine and characterized the pausing behavior of wild-type (wt) and β^R542A^ RNAP variants *in vitro*. We used a template that contains a strong T7 A1 promoter followed by the *ops* and *his* pause elements ^11^. On this template, RNAP can be halted at *ops* G5 in the absence of UTP; thus synchronized, α^32^P-labeled G5 ECs are restarted upon the addition of all NTPs. RNAP strongly pauses at the *ops* (U11) and *his* pause sites before making the run-off (RO) RNA. The wt RNAP paused at U11 with a half-life of 8 seconds, whereas βR542A substitution delayed escape ∼2.5 fold (Figure 2H), suggesting that βR542 promotes escape from the *ops* pause, in contrast to its effect at the elemental pause ^50^. The flexible sidechain of βR542 can interact with the edge of the downstream DNA (e.g., PDB IDs 5VOI, 7YPA, 8FVW) or NT DNA -1/+1 nts (e.g., PDB IDs 8EG7, 8EG8, 8EH8). Accordingly, it is conceivable that βR542 effects on pause escape may differ depending on the sequence context. In context of *ops*PEC, βR542 presumably functions as a catalyst of DNA separation by temporarily replacing base pairing with the protein-T DNA interaction.

To evaluate the role of residues that appear to stabilize the *ops*HP, we substituted arginine residues in the β lobe (R371A/R394A), β protrusion (R470A/R473A), or β’ rudder (R314A) and characterized pausing of the RNAP variants *in vitro*. We found that neither substitution had significant effects on pausing at *ops* (Figure 2H), as was also observed with base substitutions that destabilized the *ops*HP stem ^41^.

In order to assess the basis of the extension of ss DNA regions in the transcription bubble, we determined a 3.0 Å cryoEM/SPA structure of an *ops*PEC assembled on a partially non-complementary scaffold (Figure S2C, Table S2). The nc-*ops*PEC, which was virtually identical to *ops*PEC assembled on the complementary (c) scaffold (Figures S2D-S2H). Thus, the extended ss regions in the *ops*PEC form independently of the precise sequences forming the bubble and the hybrid, suggesting that this extension principally depends on the NT-DNA HP.

### RfaH recruitment proceeds *via* a hyper-paused encounter complex

In autoinhibited RfaH, the KOW^α^ masks the β’CH-binding site on NGN ^11^, yet NMR data show that autoinhibited RfaH binds RNAP near its final binding site ^42^, suggesting the existence of a transient encounter complex, *ops*PEC^Enc^. To image this complex, we used a F51C,S139C RfaH variant (RfaH^CC^) locked in the autoinhibited state by a disulfide bridge between NGN and KOW^α^ (Figure 3A). We previously showed that RfaH^CC^ is fully active under reducing conditions ^11^, and far-UV CD and 2D [^1^H,^15^N]-HSQC spectra confirmed that RfaH^CC^ adopted the expected states under non-reducing and reducing conditions, while analytical gel filtration ensured the homogeneity of the sample (Figure S3).

**Figure 3.**
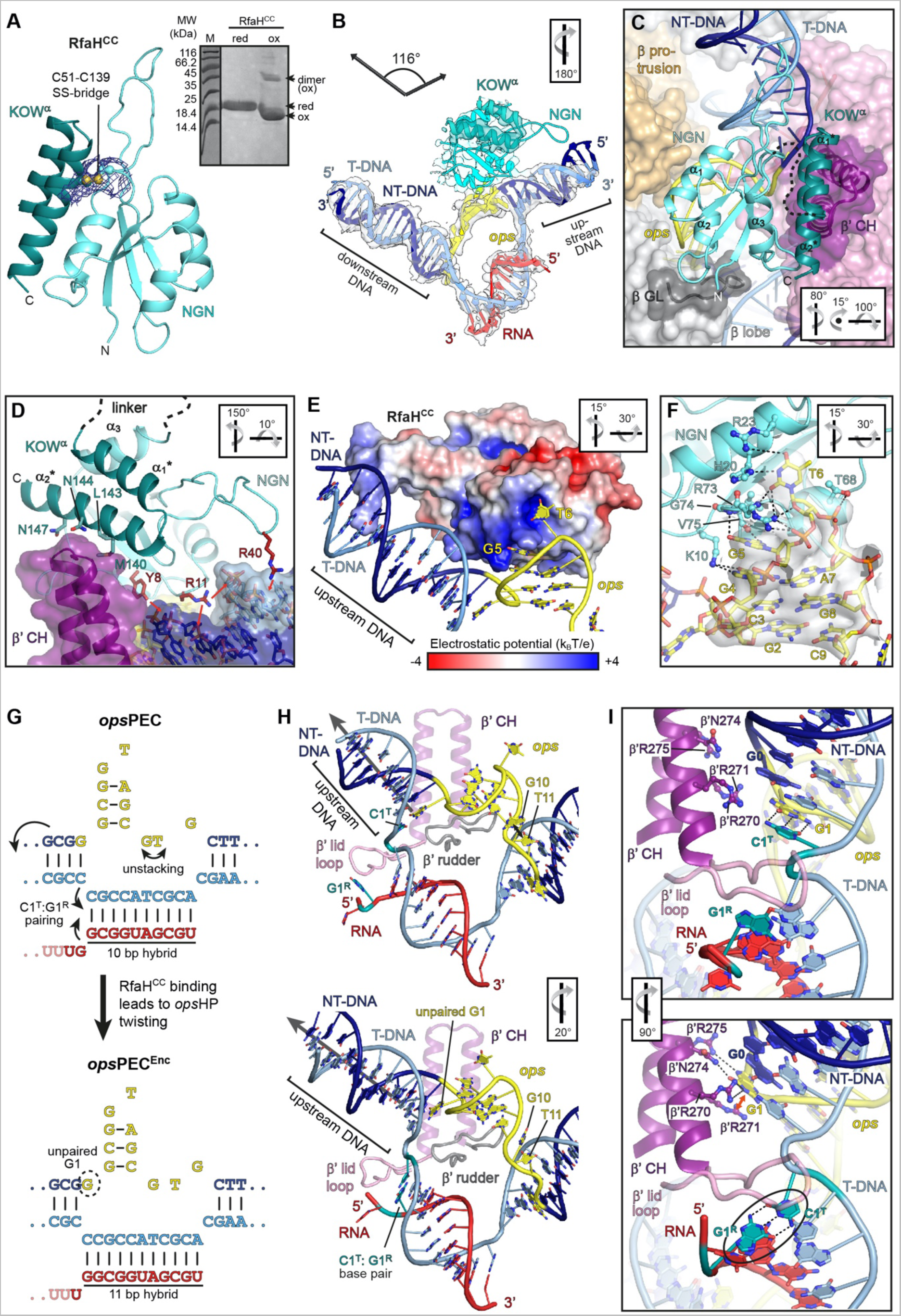
RfaH recruitment proceeds *via* an encounter complex. (A) C51-C139 bridge (balls and sticks, with cryoEM density as blue mesh) prevents domain separation in RfaH^CC^ . SDS-PAGE analysis of RfaH^CC^ in reducing (red) or non-reducing (ox) conditions. Under oxidizing conditions, the intra-molecular bridging leads to faster migration, and intermolecular bridging gives rise to dimers. (B) RfaH^CC^ bound to the nucleic acid scaffold shown as cartoon with the associated cryoEM densities (transparent surface). (C) RfaH^CC^ (cyan/mint, with several α-helices labelled and the interdomain linker shown as a dashed line) is wedged between the β’CH and *ops*HP. RNAP is shown as transparent surface, with relevant elements as color-coded cartoons. (D) (D) The β’CH act as a wedge to induce RfaH domain separation. In RfaH^CC^ and β’CH, side chains of interacting residues are shown as sticks. Nucleic acids are represented as transparent surface/sticks. RfaH^CC^ residues forming polar interactions (red arrows) with the upstream DNA are in salmon. (E) RfaH^CC^ binds *ops* and upstream DNA *via* positively charged patches. The electrostatic potential of RfaH^CC^ is mapped on its molecular surface. (F) Side chains of NGN residues contacting *ops* are depicted as sticks (salmon). The *ops* DNA is shown as sticks along with the cryoEM density of the *ops*HP (transparent surface). H-bonds and electrostatic interactions are indicated by dashed lines. (G – I) Upon binding of RfaH^CC^, *ops*PEC (top) rearranges into the hyper-paused *ops*PEC^Enc^ (bottom). (G) A summary of nucleic acid changes. (H) Nucleic acids (cartoon/stick) and β’ lid loop, β’ rudder loop and β’CH (cartoon). C1^T^ and G1^R^ form an 11^th^ bp in *ops*PEC^Enc^ (cyan). The upstream DNA vectors are indicated by arrows. (I) A close-up view of the upstream fork junction. Selected β’CH side chains are shown as sticks; polar (dashed lines) and stacking (red arrow) interactions are indicated. In panels B, C, D, E, F, H and I, orientations are relative to the standard view (Figure 2A).

We mixed RfaH^CC^ with the assembled *ops*PEC under non-reducing conditions and determined the structure of the ensuing *ops*PEC^Enc^ by cryoEM/SPA at 2.6 Å resolution (Figure 1A, Table S1). Except for the β’ zinc binding domain (ZBD), which could not be reliably modeled, most RNAP elements and the nucleic acid regions were well defined in the cryoEM reconstruction. In addition, there was a clear density for RfaH^CC^.

The *ops*PEC^Enc^ is swiveled, albeit less strongly than *ops*PEC (4.9 ° vs. 5.8 °, Figure S4A), with an unfolded TL and an open SI3. Correlating with reduced swiveling, the β’CH are slightly displaced from the β lobe/protrusion, and the angle between upstream and downstream DNA duplexes is increased compared to *ops*PEC (∼116 °) but remains smaller than in the canonical EC (Figure 3B). Therefore, sufficient space exists between the β lobe/protrusion and β’CH for RfaH^CC^ NGN to bind the *ops*HP and RNAP (Figures 3B and 3C). However, the KOW^α^-NGN interaction prevents full accommodation of RfaH as observed in RfaH-*ops*EC ^12^.

In RfaH-*ops*EC ^12^, RfaH is bound in an open conformation; NGN is positioned across the RNAP main channel, with helices α1 and α2 contacting the β protrusion and β lobe, respectively, and the opposite open flank of the central β-sheet and helix α3 contacting the β’CH. In *ops*PEC^Enc^, the NGN packs with helix α2 against the β lobe similar to RfaH-*ops*EC, whereas the NGN is rotated about the α2 axis towards the β protrusion (Figures 3C, S4B and S4C). NGN helix α1 is moved closer to, and interacts more intimately with, the β protrusion, while on the other flank, the loop preceding NGN helix α3 is displaced from the β lobe and instead interacts with the β’ clamp region neighboring the β’CH. Helix α3 is thereby moved away from the β’CH, and its rough position is instead occupied by the α2* helix of KOW^α^ (*, secondary structure elements in KOW; Figures 3C, 3D and S4B).

G5 and T6 in the *ops*HP loop serve as crucial anchors during RfaH recruitment ^41^. In full agreement, in *ops*PEC^Enc^, the *ops*HP stem has the same base pairing pattern as in *ops*PEC and its tip is engaged by NGN (Figures 3E, 3F, S4C). The NGN-*ops*HP interaction closely resembles the crystal structure of the RfaH-*ops* binary complex ^41^; *i.e.*, the *ops*HP is bound at NGN opposite KOW^α^, T6 is inserted into a positively charged pocket of NGN, and G5 packs against a neighboring surface (Figures 3D and S4D). Nucleobase-specific interactions between NGN and the *ops*HP loop (Figure 3E) show that RfaH reads out the NT-DNA sequence already in *ops*PEC^Enc^. T6 not only contacts NGN but it also stacks on βY62 (β protrusion; Figure S4C), an interaction not observed in RfaH-*ops*EC ^12^. Additionally, the same positively charged RNAP side chains as in *ops*PEC contact the *ops*HP stem, but at different chemical functionalities or nts. As a result, T6 is snuggly sandwiched between NGN and RNAP, and the entire *ops*HP is tightly restrained, as indicated by its well-defined density (Figures 3B and 3F).

Upon NGN grabbing a hold of the *ops*HP loop, the *ops*HP is repositioned, with profound consequences for the paused state. Docked RfaH^CC^ twists the *ops*HP and redirects its tip towards the upstream DNA, while G10 unstacks from T11, which retains stacking interactions with βW183 (Figures 3G, 3H and S5C). Furthermore, Y8 and R11 in the loop preceding helix α1 of RfaH^CC^ contact the sugar-phosphate backbone of upstream DNA on the major groove side (Figure 3D); based on these DNA “anchors”, the proximal T-DNA branch of the upstream duplex is pulled against the β’ rudder as RfaH^CC^ engages the *ops*HP (Figure 3H). The β’ rudder thereby acts like a strand separator, displacing C1^T^ from G1 to melt an additional upstream DNA bp (Figures 3G-3I). A β-hairpin loop (HL; M32-L50) of NGN is well-defined in *ops*PEC^Enc^, and R40 at the HL tip contacts the more distal upstream DNA backbone (Figures 3D and 3E). In *ops*EC-RfaH, the HL is disordered, possibly due to the short upstream DNA duplex employed ^12^.

Upstream DNA melting leaves G1 unpaired, and its nucleobase stacks with the neighboring upstream nt (G0); the β’CH residues (β’R271, β’N274) stabilize G1 and the new most proximal bp of the upstream DNA duplex (β’R270). Most importantly, upon RfaH^CC^-induced repositioning of the *ops*HP and upstream DNA melting, the liberated C1^T^ is paired with G1^R^, extending the RNA:DNA hybrid to 11 bps (Figures 3G-3I). As in *ops*PEC, β’L255 and β’R259 (β’ lid) cap the upstream edge of the hybrid, leading to its further compression, an increase in diameter by ∼1 Å, and, thus, a more A-like conformation compared to the 10-bp hybrid in *ops*PEC (Figures 3H and 3I). The hybrid remains pre-translocated and its downstream end matches the conformation observed in *ops*PEC, but density for the most downstream bp is less well defined (Figure 3B). The hybrid is thus pushed backward due to the destabilization of its downstream end, further counteracting translocation. Thus, pausing not only persists but is reinforced during initial RfaH recruitment, changes that might be required to allow RfaH activation and full accommodation.

We wondered how strongly the initial docking of autoinhibited RfaH onto the *ops*HP drives upstream DNA melting and, consequently, hybrid expansion. Therefore, we assembled an EC on a scaffold in which a WC bp in the upstream DNA would have to be disrupted and formation of a non-WC C:U pair would have to be “forced” to generate the 11-bp hybrid (Figure S5). The 3.1 Å cryoEM/SPA-based structure revealed that the ensuing RfaH^CC^-bound complex was virtually identical to *ops*PEC^Enc^. Strikingly, even though the C:U bp in the hybrid does not energetically fully compensate for the lost bp in the upstream DNA, the hybrid was extended to 11 bp (Figures S5E-S5G). We conclude that docking of RfaH provides a strong driving force for upstream DNA melting and hybrid expansion, leading to a hyper-paused *ops*PEC^Enc^.

The overall conformation of RfaH^CC^ in *ops*PEC^Enc^ closely resembles the structure of isolated, autoinhibited RfaH ^11^, but embedding of RfaH^CC^ between the β lobe, β protrusion and β’CH leads to a slight displacement of KOW^α^ relative to NGN (Figures 3D, S4B, S4E). While full displacement of KOW^α^ is prevented by the disulfide bridge, the β’CH tip acts like a wedge that starts to insert between NGN and KOW^α^ (Figures 3D and S4B). Concomitantly, KOW^α^ helices α1* and α2* are unwound by two N-terminal turns and one C-terminal turn, respectively (Figure S4E). Thus, upon initial docking to *ops*PEC, autoinhibited RfaH takes a handle of the *ops*HP loop to pull its own KOW^α^ against the β’CH, generating steric conflicts that prime KOW^α^ dissociation and subsequent refolding, and thus RfaH activation.

### The hyper-paused state persists after full accommodation of RfaH

To follow the complete RfaH recruitment and activation, we determined cryoEM/SPA structures of *ops*PECs assembled on c- and nc-scaffolds with RfaH^wt^ (Figures 1, 4A, S6; Tables S1 and S2). To facilitate possible conformational changes, we incubated the samples at 37 °C for 10 minutes before vitrification and imaging. CryoEM reconstructions followed by 3D variability analysis (3DVA) revealed two nearly identical states (Figure 4A). We focus on complex 1 assembled on the c-scaffold in which both RfaH domains are visible (Figure 4A, left; thereafter designated as *ops*PEC^Rec^) and discuss the alternative state below.

**Figure 4.**
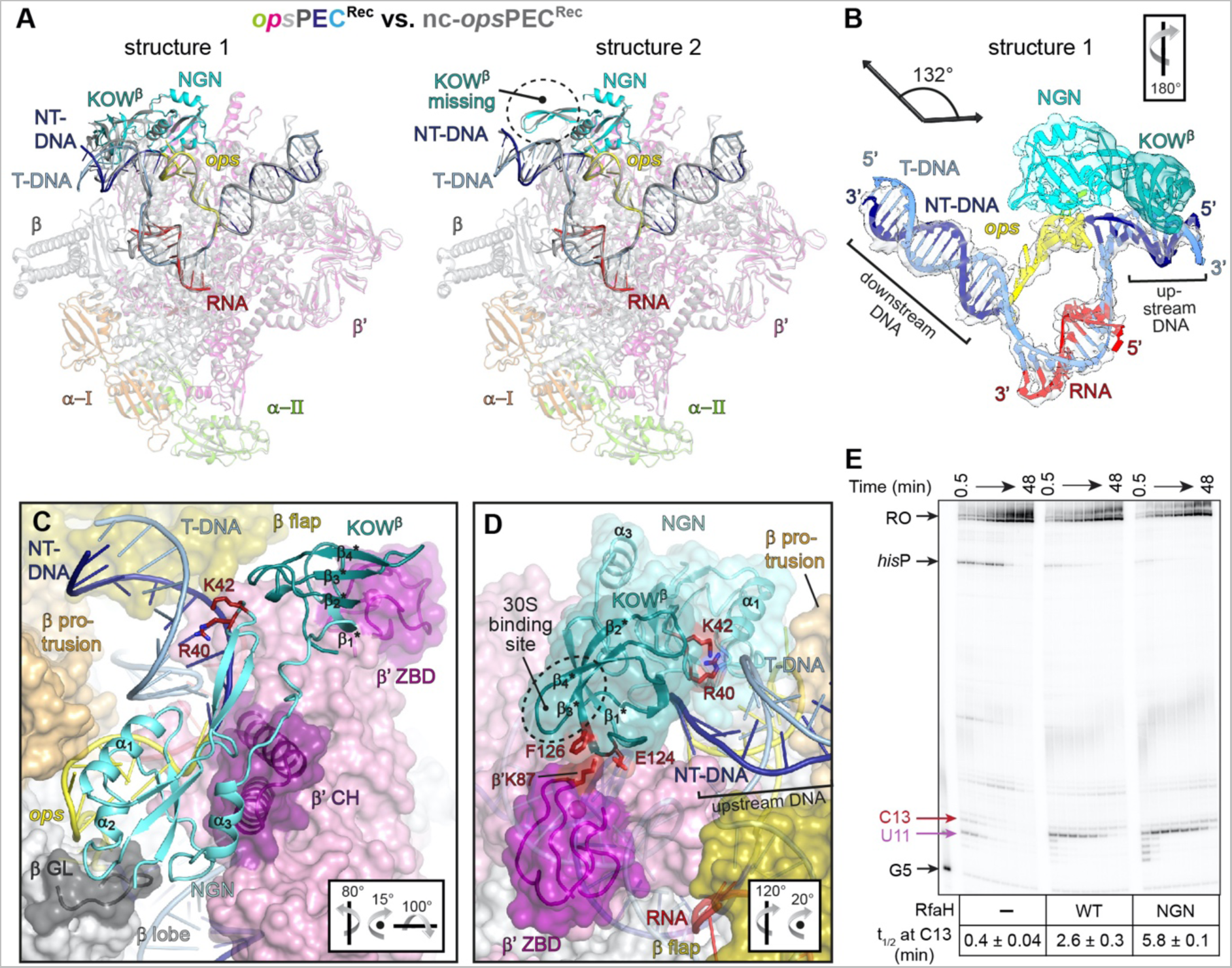
Recruited RfaH is activated and keeps *ops*PEC in the hyper-paused state. (A) Superposition of *ops*PEC^Rec^ (in colour) and nc-*ops*PEC^Rec^ (grey). 3DVA of *ops*PEC^Rec^ cryoEM maps revealed two extrema, structure 1 (left) with KOW^β^ bound to the β’ZBD and poorly ordered *ops*HP DNA (the latter only observed for nc-*ops*PEC^Rec^) and structure 2 (right) exhibiting no cryoEM density of KOW^β^ but well-defined *ops*HP density. (B) *ops*PEC^Rec^ structure 1. The nucleic acids and RfaH are shown as coloured cartoons along with their corresponding cryoEM density (transparent surface). The helix axes vectors of up- and downstream DNA and the angle between them are shown on top. (C and D) Positioning of RfaH within *ops*PEC^Rec^. RNAP is shown as transparent surface, selected structural segments are depicted as cartoons. RfaH secondary structure elements are labelled. In (D), KOW^β^ contacts to the β’ZBD and upstream DNA; side chains of interacting residues are displayed as red sticks. In panels B-D, the orientation is relative to panel A. (A) (E) Deletion of KOW augments RfaH-induced pausing at C13. Halted radiolabelled G5 ECs were chased in the absence of RfaH or in the presence full-length RfaH or NGN, as described in STAR Methods. Samples withdrawn at 0.5, 1, 2, 4, 6, 12, 24, and 48 min were analyzed on a urea-acrylamide gel. The positions of *ops* G5, U11 and C13, *his*P, and run-off (RO) RNAs are indicated with arrows. The half-life (t_1/2_) of pausing at C13 is presented as mean ± SD, n=3.

RfaH is bound in its activated state (Figure 4B), with NGN fully embedded between the β lobe, β protrusion and β’CH, as observed in RfaH-*ops*EC ^12^. As compared to *ops*PEC^Enc^, NGN is rotated about the axis of helix α2, so that α1 is slightly displaced from the β protrusion, while α3 moves to contact the β’CH (Figure 4C). In contrast to RfaH-*ops*EC, *ops*PEC^Rec^ retains a swiveled conformation, albeit with a reduced swiveling angle of 3.3 ° as compared to *ops*PEC^Enc^ (4.9 °). Thus, RfaH binding *per se* does not prevent swiveling as previously suggested ^12^.

The *ops*HP conformation and position are essentially unaltered compared to *ops*PEC^Enc^ and very similar to RfaH-*ops*EC (Figures 4B vs 3B). However, the loop preceding RfaH helix α1 is moved closer to the upstream DNA and engages in more intimate interactions with the sugar-phosphate backbone on the major groove side; the HL-upstream DNA contacts are also maintained, albeit to a more proximal region of the upstream duplex (Figure 4C). We observed clear density for the refolded KOW^β^, which is positioned on top of the HL and contacts the β’ZBD, which becomes ordered (Figures 4B and 4C). KOW^β^ residues E124 and F126 in the loop connecting β1* and β2* sandwich the β’ZBD K87 and the upstream DNA is displaced towards the β protrusion and β flap (Figures 4C and 4D). In RfaH-*ops*EC, KOW^β^ was observed in a similar location ^12^, also interacting with the β flap tip helix (FTH). In *ops*PEC^Rec^, in contrast, the βFTH remains disordered, possibly because the longer upstream DNA we employed prevents close approach to KOW^β^. Notably, the binding site for S10 is exposed in KOW^β^.

In *ops*PEC^Rec^ the upstream duplex is pushed against the β’ rudder, maintaining the additional melted bp, and the hybrid is compressed and pre-translocated. The same configuration is also adopted in *ops*PEC^Rec^ assembled on the nc-scaffold, showing that the strong driving force for upstream DNA melting and hybrid expansion is maintained in *ops*PEC^Rec^. In summary, *ops*PEC^Rec^ remains hyper-paused, with KOW^β^ poised to engage a ribosome to form an RfaH-bridged expressome.

### Refolding landscape of RfaH upon recruitment to *ops*PEC

Ribosomal interactions with its KOW^β^ element are critical for the cellular function of RfaH ^43^. Thus, the KOW fold-switch is the final step in RfaH activation. Our structures capture the autoinhibited and activated states of RfaH-bound *ops*PECs but provide no information about their interconversion. Molecular dynamics (MD) simulations have been used to interrogate the KOW switch ^51^, but these simulations were done with the isolated RfaH or KOW. To explore the fold-switch of RfaH bound to *ops*PEC, we generated an all-atom dual-basin structure-based model (SBM), such as those employed to study massive structural transitions of influenza hemagglutinin ^52^ and SARS-CoV-2 spike protein ^53^. In 500 independent runs performed using a dual-basin SBM created based on *ops*PEC^Enc^ and *ops*PEC^Rec^, KOW underwent a complete α-to-β fold-switch.

Figure 5 shows the refolding landscape of RfaH projected onto the fraction of interdomain (ID) contacts (Q_ID_) and the difference in the fraction of native contacts formed with respect to either KOW^α^ or KOW^β^ (Q_diff_). To easily visualize KOW^β^ when Q_ID_ = 0, we employed the distance between the NGN and KOW instead. RfaH fold-switch (Figure 5A) requires that at least 60% of the ID contacts are broken (Figure 5B). Following the fraction of formed native contacts for each KOW fold (Figure 5C) reveals that RfaH refolding is rugged, with at least four intermediate states, I1-I4. I1 and I2 are connected to the KOW^α^ basin, whereas I3 and I4 have a higher fraction of KOW^β^-like native contacts (Figure 5D). Refolding of RfaH starts by the loss of native contacts at the N-terminus of the KOW α1* helix (residues 117-122, I1), followed by unwinding of the end of KOW α2* helix (residues 148-155; I2) (Figures 5D and 5E). Notably, this occurs as observed in the *ops*PEC^Enc^ cryoEM structure (Figures 3D and S4E) even though the helical content of these regions was restored using homology modelling, and is in agreement with hydrogen-deuterium exchange mass spectrometry data showing that the hairpin tip is more stable than the termini of the helices ^54^. Most ID interactions, except for those between NGN 85-100 and KOW 114-126/150-162 from the ends of KOW α1*/α2* helices, persist through I2.

**Figure 5.**
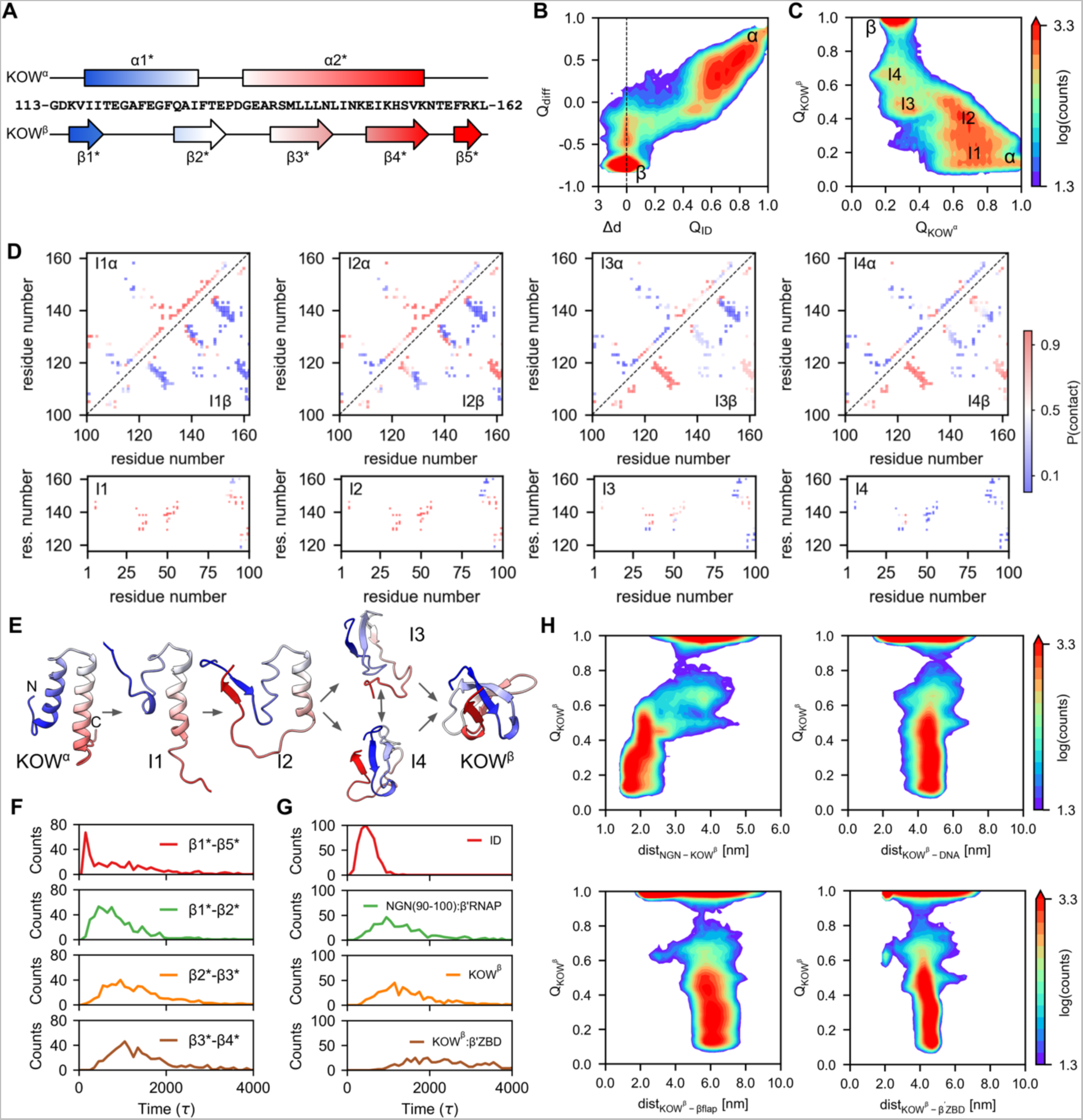
Refolding landscape of RfaH. **(A)** Secondary structure topology of the KOW^α^ (top) and KOW^β^ (bottom). Helices are represented as rectangles and strands as arrows. (B) Refolding landscape of RfaH projected onto *Q*_ID_ (fraction of ID contacts), Δ*d* (distance between domains with respect to the distance in the active state, in nm) and *Q*_diff_ (difference in native contacts between the KOW^α^ and KOW^β^). The color scheme represents the number of times each configuration is observed across all MD simulations. (C) Refolding landscape projected onto the fraction of native contacts of each KOW state (*Q*_KOW_^α^ and Q_KOW_^β^). Intermediate states are labeled. (D) Probability of native contacts belonging to either KOW^α^ (upper triangle), KOW^β^ (lower triangle) or ID contacts (bottom plots) present in each intermediate state. (E) Intermediates in the KOW refolding pathway. (F) Histograms of the number of dissociation and association events as a function of time, with domain dissociation being the first event during refolding. (G) Histograms of the number of β-strand formation events as a function of time. (H) Landscapes of the KOW refolding as a function of the distance between KOW and NGN, DNA, β flap and β’ZBD.

In I2, KOW β5* is released from NGN and interacts with β1* (average contact probability 0.9), and emergent interactions are also observed between strands β1*-β2* and β1*-β5* (Figure 5D). In I3, the probability of forming native β1*-β2* contacts (on average, 0.9) surpasses that of β1*-β5* (on average, 0.6) and β2*-β3* (on average, 0.4). These strands are formed just after completion of the α1* unwinding and the loss of most interhelical and ID interactions, except for contacts between the KOW hairpin tip and NGN (Figure 5D). Last, I4 is characterized by nearly complete unwinding of α2* (residues 137-154), the loss of almost all ID interactions, and high probability native interactions between strands β1*-β5*, β2*-β3* and β3*-β4* that will later consolidate the KOW^β^ (Figure 5E). The time course of β-strand KOW folding events (Figure 5F) follows the sequential pattern, with the early formation of the interactions between β1*-β5* (peak at 100 *τ*), followed by β1*-β2* (800 *τ*), and lastly β2*-β3* and β3*-β4* (1000 *τ*). The refolding trajectories are heterogeneous regarding the order of formation of β1*-β5* (Movie S1) or β1*-β2* (Movie S2) interactions.

Next, we analyzed the sequence of events within RfaH and between RfaH and *ops*PEC during the transition from *ops*PEC^Enc^ to *ops*PEC^Rec^. We examined the disruption of native ID contacts and the formation of native contacts for KOW^β^, contacts between the NGN α3 helix (residues 90-100) and RNAP, and contacts between KOW^β^ and the β’ZBD as a function of time. The first event enabling RfaH refolding is the breakage of 70% of ID interactions (peak at 400 *τ*, Figure 5G), in line with the evidence that the NGN-KOW contacts control RfaH metamorphosis ^55, 56^. Either concurrently or after domain dissociation (1000 *τ*), α3, which harbors a highly conserved I93 residue that stabilizes the ID interface ^57^, binds to β’CH (Movies S1 and S2). In these simulations, α3 gets locked in place by forming >75% of its native contacts with RNAP and enabling tighter binding of RfaH to *ops*PEC. Then, KOW refolds into the β-barrel (peak at 1100 *τ*). This event is not concurrent with binding to DNA, β flap or β’ZBD (Figure 5H and Movies S1 and S2).

Our simulations reveal that the KOW switch is initiated by unwinding of the ends of its α- helices, proceeds via a largely unfolded state that transiently retains some helical elements at the tip of the KOW hairpin, culminating with the formation of KOW^β^ through heterogeneous intermediate states where different β-strands come together. These findings are in agreement with our structural (Figure 3) and biophysical analyses ^54, 58^ and with MD simulations performed in the absence of RNAP ^55, 59, 60^. We conclude that the KOW transformation is independent of its interactions with the *ops*PEC.

### The *ops*PEC^Rec^ can be arrested

The *ops*PEC has been identified as a class II, or backtrack-prone, pause ^44^ and our results show that RfaH further stabilizes the paused state. How does *ops*PEC^Rec^ resume elongation? To answer this question, we subjected *ops*PEC^Rec^ after NTP addition and heating (37 °C) to cryoEM/SPA, yielding a cryoEM reconstruction at 3.0 Å resolution. The resulting complex was nearly identical to *ops*PEC^Rec^ (Figures 6A and S7A; Table S1) with a major exception: a clear additional density in the secondary channel showed that the transcript had been elongated by at least two nts followed by RNAP backtracking (Figures 6B and 6C). These findings establish that *ops*PEC^Rec^ is elongation-competent and that, following nucleotide addition, RNAP slides back, generating *ops*PEC^Back^.

**Figure 6.**
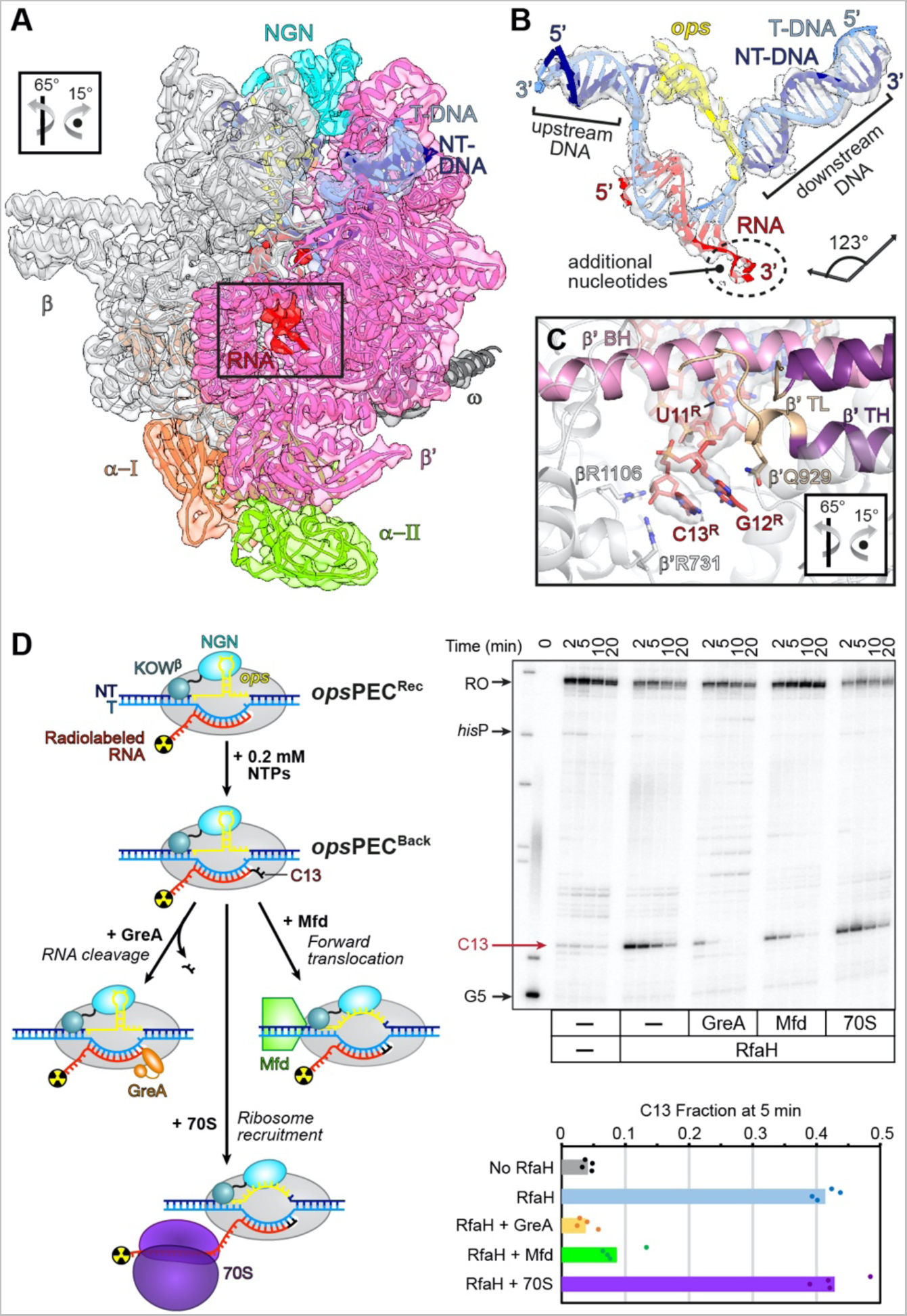
RNAP escape from *ops* is hindered by backtracking and requires auxiliary factors. **(A)** Upon nucleotide addition, *ops*PEC^Rec^ forms *ops*PEC^Back^ (map as transparent surface, model as cartoon) with two additional RNA nucleotides in the secondary channel (boxed). (B) CryoEM map and model of the *ops*PEC^Back^ nucleic acid scaffold. Additional 3’ RNA nts resulting from RNA extension and subsequent backtracking are indicated. (C) Close-up view of the region boxed in (A). RNAP, β’BH and β‘TL are shown as cartoons, RNA – as sticks along with the corresponding cryoEM density. RNAP side chains contacting the protruding RNA are shown as sticks. In A and C, the orientation is relative to Figure 2A. (D) The arrested *ops*PEC^Back^ can be rescued by either GreA-assisted RNA cleavage or Mfd-medicated forward translocation. Halted radiolabelled G5 ECs were chased with 0.2 mM NTPs in the absence or in the presence of indicated proteins, Samples withdrawn at 2, 5, 10, and 20 min were analyzed on a urea-acrylamide gel. The positions of *ops* G5 and C13, *his*P, and run-off (RO) RNAs are indicated. RNA fractions at C13 after 5 min incubation with NTPs were calculated. Raw data points (scattered dots) and the mean values are shown; n=4.

In the cell, backtracked RNAP can be rescued by Gre factors ^49^, Mfd ^61^, or the coupled ribosome ^62^. We next tested if RNAP escape from *ops* can be promoted by anti-backtracking factors *in vitro*. In agreement with our structural data, RfaH strongly delays elongation two nts downstream from the *ops* pause site: the C13 pause persists for minutes even at 0.2 mM NTPs (Figure 6D). The addition of GreA, which induces the nascent RNA cleavage in backtracked ECs ^49^, dramatically shortened the pause. A similar but less dramatic effect was observed in the presence of Mfd, a DNA translocase that pushes RNAP forward ^61^. By contrast, the 70S ribosome did not promote escape, an expected result given that the ribosome must exert force on RNAP to assist forward translocation, not just sterically block reverse translocation ^62^, and translation initiates only 50+ nts downstream from the *ops* site.

### The KOW domain contributes to pause escape

3DVA of *ops*PEC^Rec^ revealed that KOW^β^ binding at the β’ZBD was correlated with a movement of the upstream DNA towards the β protrusion and β flap (Movies S3 and S4; Figure S6). The apparent push of KOW^β^ on upstream DNA suggests that it reinforces HL-upstream DNA contacts, which are already established in *ops*PEC^Enc^ and may either support hyper-pausing by helping drive the proximal end of the upstream duplex into the β’ rudder or, alternatively, could counteract backtracking of *ops*PEC^Rec^. The removal of KOW potentiates the RfaH-induced delay at C13 (Figure 4E). While a fraction of arrested complexes may escape by releasing RfaH and reformation of the autoinhibited state in full-length RfaH, but not in the isolated NGN, this observation is consistent with the idea that, after RfaH accommodation and refolding, KOW^β^/HL act as anti-backtracking devices. A similar effect has been observed for SuhB-reinforced NusG-upstream DNA contacts in an rRNA antitermination complex ^6^.

3DVA also showed that in some complexes, density for KOW^β^ was anti-correlated with density for the *ops*HP; *i.e.*, one boundary state exhibited clear density for KOW^β^ bound at the β’ZBD but weak density for the *ops*HP and downstream *ops* nts, whereas the other lacked density for KOW^β^ but exhibited very well-defined density for the *ops*HP and downstream nts (Movies S3 and S4). Thus, in addition to dampening backtracking, KOW^β^ binding at the β’ZBD seems to facilitate pause escape by destabilizing *ops*HP-NGN interactions.

### NusA and KOW may cooperate during ribosome loading

NusA, a general elongation factor that associates with most ECs ^3^, modulates RNAP pausing, termination, and antitermination through contacts to the nascent RNA or accessory factors ^6, 45, 63–65^ and can aid in coupling transcription to translation ^20^. NusA binds to the βFTH ^45^ and is expected to associate with, and possibly trigger conformational changes in, RfaH-bound ECs. To ascertain that our conclusions would hold in the presence of NusA, we assembled a NusA-modified *ops*PEC^Rec^ and determined its structure by cryoEM/SPA at 3.2 Å resolution. The cryoEM reconstruction revealed that, except for added NusA, the ensuing complex is essentially identical to *ops*PEC^Rec^, suggesting that NusA binding *per se* does not induce conformational changes in *ops*PEC^Rec^ (Figure S7) and does not alter the KOW^β^ presentation. Thus, NusA could provide yet another platform for ribosome engagement through direct contacts to the ribosome ^20^.

## DISCUSSION

In this work, we present a series of structures that delineate co-transcriptional recruitment and activation of RfaH (Figure 7). We show that *ops*PEC is an unusual example of the consensus pause in which the *ops* element, a target of RfaH, forms a short NT-DNA hairpin, *ops*HP, exposed on the RNAP surface. Folding of the *ops*HP and its interactions with specific RNAP elements push upstream DNA into the β’ zipper and β’CH, thereby supporting RNAP swiveling, and stabilize a paused pre-translocated state with a 10-bp hybrid and extended bubble. The autoinhibited RfaH, in which the RNAP-binding site on NGN is partially masked by KOW^α^, docks near its final binding site using the *ops*HP as an anchor to form an encounter complex, in which NGN grasps and twists the *ops*HP to hyper-stabilize the PEC with an 11-bp hybrid. Next, RfaH domains dissociate; NGN takes its final position whereas KOW refolds into the β-barrel. Our structural data and MD simulations show that while binding to the *ops*PEC triggers the initial domain separation/refolding, the subsequent KOW fold-switch is spontaneous. Following refolding, KOW^β^ may form transient contacts with RNAP awaiting the arrival of the ribosome. Our work portrays a remarkably economical mechanism of deoxyriboregulation of RNAP, in which a merely 12-nt region of NT DNA directs major conformational changes in the transcription machinery that trigger further modulation via a metamorphic accessory factor, ultimately supporting synthesis of vital proteins.

**Figure 7.**
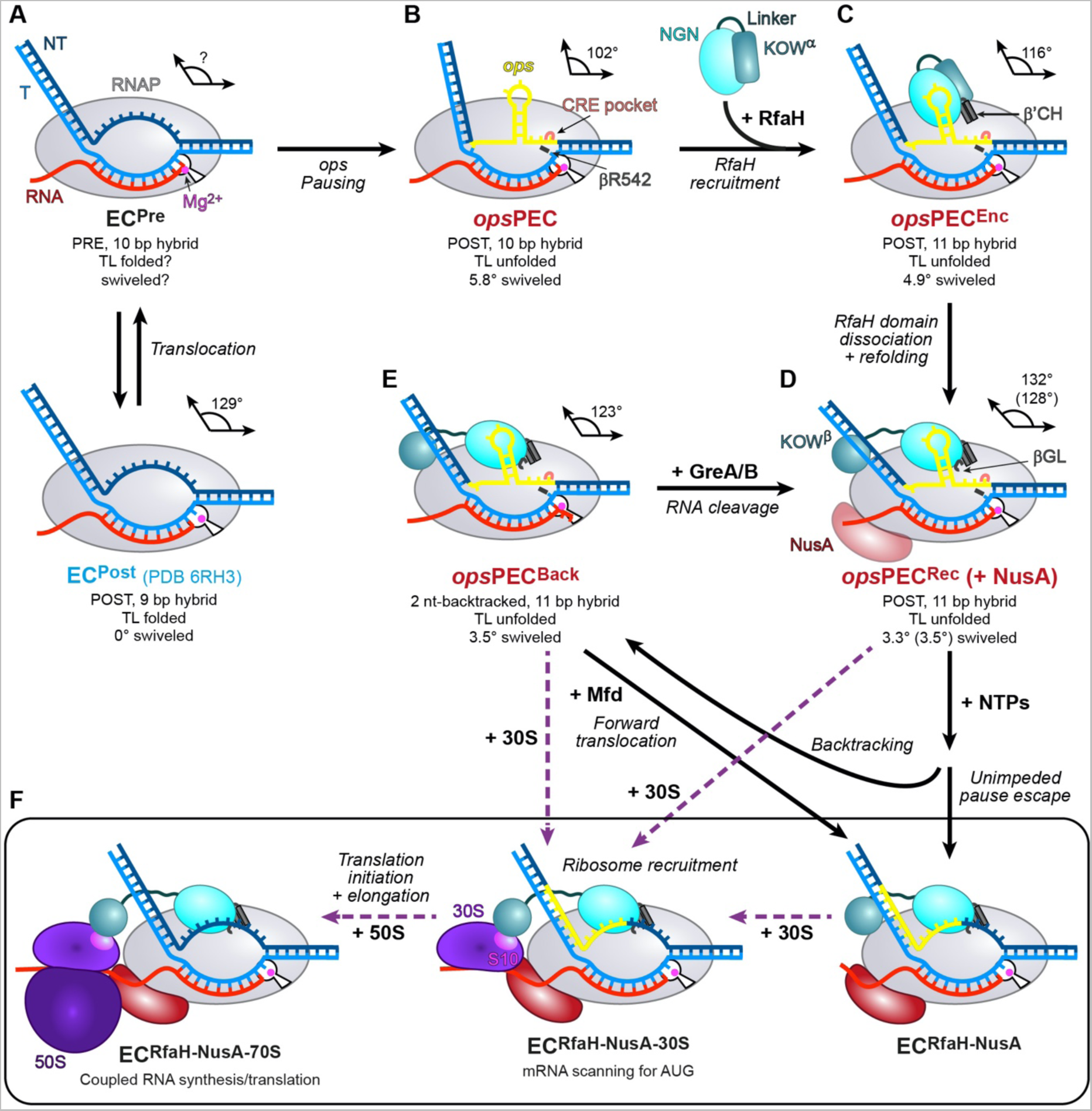
Model of transcription regulation by RfaH. Schematics of relevant ECs/PECs. Complexes with previously known structures are labeled in blue, complexes with structures determined in this study in red, and hypothetical complexes in black. See main text for details.

### The mechanism of pausing at the *ops* site

During processive RNA synthesis, RNAP adds nucleotides one by one to the 3’-end of the growing RNA. Following nucleotide addition, the NTP in the active site turns into the RNA 3’-end and the 9-bp RNA:DNA hybrid converts into the 10-bp hybrid. RNAP must then translocate by 1 nt, melting 1 bp of the downstream DNA to place the next T-DNA base in the active site, ready to accept the incoming NTP (Figure 7A); the most upstream bp of the hybrid is broken and T- and NT-stands reanneal ^66^. A failure to translocate leads to pausing, a key regulatory mechanism used to (i) control the speed of RNA synthesis; (ii) coordinate transcription with other processes such as translation; (iii) recruit regulatory factors; *etc.* ^67^.

Pausing at *ops* is a prerequisite for RfaH recruitment ^11^. The *ops* element stands out as one of the strongest pauses in *E. coli* ^47^, as could be expected because a failure of RfaH engagement compromises the cell wall integrity ^68^, and an exemplar of a consensus pause sequence identified by genome-wide analysis. ^47, 48^ The *ops*PEC structure (Figure 7B) reveals a unique geometry of the nucleic acid scaffold, a canonical 10-bp hybrid but 11-nt long ss NT DNA region instead of 10 in the typical pre-translocated EC ^49^. The NT DNA forms the *ops*HP, which comprises a 3-bp stem and is stabilized by a multitude of positively-charged RNAP residues (Figure 2D); the loop residues G5 and T6 are flipped-out. The two downstream nts, G10 and T11, stack on each other and are further stabilized by βW183, whereas G12 is flipped into the CRE pocket, leaving the C12^T^ base free to contact βR542 (Figure 2G).

Two other features of *ops*PEC are expected to contribute to pause strength ^67^. The TL is trapped in an unfolded conformation by salt bridges between the TL β’R933 and the fork loop βE546/D549 (Figure 2C). The swivel module rotates by 5.8° (Figure 2B) and is stabilized by *ops*HP interactions with the β lobe. Recently, it has been suggested that NusG-dependent pausing in Gram-positive bacteria relies on a similar mechanism wherein the NusG-bound NT DNA is placed in a cleft between NusG and the β lobe, blocking RNAP return to the non-swiveled state ^69^. Strikingly, through contacts to RNAP, the *ops*HP is able to lock the swivel module in the absence of additional regulatory inputs.

### Sequence-structure relationships in PECs

Comparison of *ops*PEC with structurally-characterized PECs, *his*PEC ^45, 70^, *his*-ePEC ^71^ and con-ePEC ^71^, offers insights into the mechanism of pausing and the contributions of the individual sequence determinants. Con-ePEC is paused on a synthetic sequence GG-CAUAGUUG-CG based on the pause consensus (^-12^GG-NNNNNNNN-YG^+1^) ^47, 48^; GG and YG motifs are known as upstream and downstream fork junction (UFJ and DFJ) determinants, respectively. The consensus within the 8N region is weak and, in the first approximation, the con-ePEC structure reveals contributions of DFJ and UFJ to pausing.

*ops*PEC (GG-CGGUAGCG-UG) and *his*PEC (upstream RNA hairpin followed by CG-AUGUGUGC-UG) form on naturally occurring regulatory pause elements featuring nucleic acid hairpins. In *ops*PEC, the hairpin forms in the NT DNA and is encoded by the 8N sequence, and perfect DFJ and UFJ are present. In *his*PEC, a nascent RNA hairpin forms in the exit channel. *his*PEC and *his*-ePEC, its hairpin-less derivative, have perfect DFJ and a partial UFJ.

The common feature of all PECs is failure to translocate and load the acceptor T-DNA base into the active site. *ops*PEC and con-ePEC are pre-translocated, *his*-ePEC equilibrates between pre- and half-translocated states (only RNA translocates), and *his*PEC is half-translocated. The asynchronous (half) translocation leads to tilting of the nucleobases of the RNA:DNA hybrid relative to the helical axis. Since incomplete translocation is characteristic for all pauses, the DFJ and UFJ are likely mainly responsible for the translocation block.

The synthetic con-ePEC and *his*-ePEC equilibrate between several states with open and closed active sites ^71^. In contrast, natural *ops*PEC and *his*PEC are each represented by a single state with an open active site, likely because both are strongly stabilized in a swiveled conformation by nucleic acid hairpins, and swiveling is incompatible with the folded TL ^70^. Noteworthy, a fraction of *his*-ePEC is also swiveled despite lacking the stabilizing hairpin. The causative relationships between hybrid tilting and swiveling were hitherto difficult to delineate. Considering that *ops*PEC is swiveled but not tilted suggests that tilting of hisPEC/ePEC is mediated by the 8N sequence. Indeed, AUGUGUGC consists entirely of purine-pyrimidine steps, has diminished intra-strand base stacking, and is expected to favor tilting of nucleobases to enhance the inter-strand stacking of purines. Interestingly, while both *ops*PEC and *his*PEC are swiveled, the swiveling is seemingly attributable not to the UFJ and DFJ determinants, but to pause-specific 8N regions, the *ops*HP and the tilted hybrid, respectively. In the latter case, swiveling is further stabilized by the RNA hairpin.

A notable feature of the con-ePEC is an overextended 11-bp RNA:DNA hybrid, which forms only upon active site opening and is not linked to swiveling – all states of con-ePEC are not swiveled. *ops*PEC also forms an 11-bp hybrid, but only upon RfaH binding. It is tempting to speculate that *ops*HP inhibits the formation of the overextended hybrid, whereas RfaH binding repositions the *ops*HP and unleashes the intrinsic potential of *ops*PEC to form such a state. Hybrid overextension is a step backward and may conceivably delay translocation, thereby strengthening the pause, but the underlying sequence determinants are difficult to pinpoint unambiguously. One possibility is that the DFJ and UFJ determinants are sufficient to cause hybrid overextension, and the inability of *his*PEC/ePEC to form such a state can then be attributed to hybrid tilting, the partial UFJ, or both. Another possibility is that specific determinants within the 8N sequence are the reason for hybrid overextension: con-ePEC and *ops*PEC, but not *his*PEC/ePEC, share G at -2 and C at -9.

### The role of *ops* DNA hairpin in recruitment of RfaH

Our structural analysis reveals that the *ops*HP forms in the absence of RfaH and recruits the autoinhibited RfaH^CC^ through specific contacts to G5 and T6 residues in the hairpin loop (Figure 3F). The *ops* pause is a composite element where UFJ and DFJ mediate pausing and nucleobases in the hairpin loop make specific contacts to RfaH ^41^. Many potential *ops*-like hairpins, which differ only in the loop residues and are thus expected to induce pausing, are encoded in the NT strand of MG1655 operons; RfaH is recruited only to *ops* sites that have a T at position 6 ^38^, but these comprise fewer than half of *ops*-like sequences. Do NT-strand hairpins that have A, G or C in the loop recruit, or perhaps exclude, other modulators of transcription elongation? Housekeeping NusGs from *Bacillus subtilis* ^69^ and *Mycobacterium tuberculosis* ^72^ interact with the NT DNA and *B. subtilis* NusG displays a preference for T-tracks ^73^. *E. coli* NusG makes no contacts to DNA ^12^, but could conceivably be excluded from the EC by NT-DNA structures. By contrast, RfaH orthologs must avoid recruitment to “wrong” sites, and the NT DNA readout provides means for the exquisite selectivity.

Here we show how RfaH, still in the autoinhibited state, sets the stage for the final NGN placement, with three distinct elements making contacts to the *ops*HP, β’CH and β gate loop. In the encounter complex (Figure 7C), NGN cannot be fully accommodated between the β lobe, β protrusion, and β’CH, and RNAP swiveling is required to fit the additional bulk of KOW^α^ between these RNAP elements. While anchored at the upstream DNA duplex, NGN stabilizes, twists and repositions the *ops*HP, leading to melting of one bp in the upstream DNA and inclusion of the melted T-strand nt into the overextended 11-bp hybrid, which counteracts translocation and stabilizes the pause. RfaH^CC^ clamps the NT strand between itself and the β lobe, further hindering return to the non-swiveled state and thus strengthening the pause. Contacts with *ops*HP position NGN near the β’CH tip, which starts to wedge between the RfaH domains, initiating displacement of KOW^α^ (Figure 3D). Consequently, RfaH “pulls” itself towards its final binding sites, driving the wedge further and further between KOW and NGN until they dissociate completely. Thus, the *ops*HP loop has two functions: (i) sequence-specific recognition by RfaH and (ii) anchoring RfaH to initiate its activation.

### RfaH accommodation and activation

The *ops*PEC^Rec^ structure shows that, once unmasked, the NGN helix α3 packs against the β’CH, the principal RfaH-binding site on RNAP. The *ops*HP:NGN interactions are identical to those in *ops*PEC^Enc^, while the region around T73 contacts the βGL, the second RNAP-binding site (Figure 7D). The NGN-RNAP interactions account for the remarkable stability of RfaH-EC contacts throughout transcription and for the anti-pausing activity of RfaH *in vitro* ^11, 12, 37^ but make only a small contribution to its overall effect on gene expression ^14^. The KOW binding to ribosome, thought to mediate both the initial ribosome loading and transcription-translation coupling, is critical for RfaH activity and is, in turn, dependent on refolding of the liberated KOW^α^ into a NusG-like β-barrel that creates a contact surface for S10 ^43^.

KOW refolds spontaneously when freed from NGN, as demonstrated by NMR spectroscopy ^41, 43^, but the fold-switch could be altered in the context of the *ops*PEC, due to KOW-EC contacts revealed by structures. Our MD simulations reveal that refolding starts from the ends of the α-helical KOW hairpin (Figure 5) consistent with its partial unfolding in the *ops*PEC^Enc^. Once the NGN α3 helix is released and gets locked in place, KOW refolds into the five-stranded β-barrel (Movie S2), largely as observed with RfaH in isolation ^51^. We conclude that all the information required for KOW transformation is encoded in its primary sequence, and that its contacts to DNA and RNAP are established following the fold-switch.

### Pause escape and recruitment of the ribosome

RfaH hyper-stabilizes the *ops* pause, an effect opposite to its pause-suppressing activity at any other site ^40^. We show that *ops*PEC^Rec^ can extend the RNA but, unable to break NGN-*ops*HP interactions, backtracks to the original position (Figure 7E). This is reminiscent of PECs stabilized by the initiation σ-factor, which can extend RNA and the bubble, scrunching both DNA strands to maintain contacts to the -10-like NT DNA ^74^. The scrunched σ-PECs can escape to elongation, breaking these contacts, or backtrack to the starting position ^74^.

It is hypothesized that the energy stored in the scrunched strands is used to break the σ-DNA contacts to overcome the pause ^74^. We do not observe scrunching in *ops*PEC^Rec^ or *ops*PEC^Back^, but the transition to *ops*PEC^Back^ must be accompanied by at least two translocation steps along the DNA and likely requires scrunching, as proposed earlier ^75^. The energy stored in the compressed bubble/hybrid and hypothetical scrunched states could be used to break the NGN/NT contacts and melt the *ops*HP to overcome the pause.

Rather than being an unavoidable consequence of overly stable protein-DNA contacts, pausing mediated by specific recognition of NT DNA may facilitate recruitment of protein partners to the EC. The phage λ Q antiterminator loads onto RNAP at a σ-dependent promoter-proximal pause through direct contacts to σ ^76^. We hypothesize that a ribosome is recruited at the *ops* site through direct contacts to KOW^β^, and that backtracking extends the time window for this recruitment. By bridging RNAP and ribosome, RfaH will promote ribosome scanning the mRNA for a start codon, where the translation initiation complex assembles, and coupling thereafter (Figure 7F). Failure to load the ribosome abolishes the expression of RfaH-controlled genes ^32^. It remains to be determined which form of the ribosome is recruited by RfaH and when, and whether RNA is required during the initial loading.

### Concluding remarks

Collectively, our results show that the information encoded in the *ops* and RfaH sequences fully controls every step in the RfaH cycle – from RNAP pausing and RfaH loading (*ops*) to RfaH activation and finally ribosome recruitment (RfaH). The KOW transformation that activates RfaH, initially thought to be unique to that protein, has recently been discovered to be ancient and ubiquitous ^77^, arguing that our insights into the mechanisms of recruitment and metamorphosis of RfaH would be applicable to NusG homologs across all life.

Here, we deciphered the molecular details that control the fold-switch in RfaH, providing insights into other fold-switch mechanisms. Recent estimates suggest that ∼4 % of structures deposited in the PDB may belong to fold-switching proteins ^78^, a figure that almost certainly understates the prevalence and significance of this phenomenon, given the bias for a single, lowest-energy structure in the most commonly used structural methods. It is worth noting that despite its relatively small size (50 residues in the KOW domain and 162 residues overall), RfaH nonetheless undergoes the most drastic fold-switch possible – from all-α to all-β. Having shown here that the sequence of RfaH itself is responsible for this transformation, we have definitively established the suitability of RfaH as an eminently tractable (both experimentally and computationally) model system for investigating the molecular determinants of protein fold-switching.

## LIMITATIONS OF THE STUDY

Our current cryo-EM reconstructions capture snapshots of key intermediates in the RfaH activation pathway but provide no information on transitions between these states. For example, we did not observe scrunching, which we presume accompanies nucleotide addition when translocation is hindered by RfaH contacts to *ops*HP. The encounter complex was obtained with RfaH^CC^ in which the domains are locked by a disulfide bridge; although we have shown that RfaH^CC^ behaves as the unmodified RfaH under reducing conditions, we cannot exclude minor differences in recruitment. The ribosome loading, a critical step in RfaH function, is a black box. Following RfaH recruitment and activation, RNAP translocation, and accompanying structural changes in the EC in real time would be required to investigate this possibility, using single-molecule approaches to account for the system heterogeneity. Finally, *in vitro* experiments have limitations that can only be overcome by studying the process *in vivo*, which may be possible due to advances in super-resolution microscopy.

## ACKNOWLEDGMENTS

This work was supported by the Deutsche Forschungsgemeinschaft (WA 1126/11-1, project number 433623608, to M.C.W.; INST 130/1064-1 FUGG to Freie Universität Berlin; Ro617/21-1 to P. Rösch), National Institutes of Health (GM067153 to I.A.), the National Agency for Research and Development (ANID) through Millennium Science Initiative Program (ICN17_022), Fondo de Desarrollo Científico y Tecnológico (FONDECYT 1201684 to C.A.R.-S) and the Academy of Finland (341962 to G.A.B.). J.G-H. was supported by an ANID Doctoral Scholarship (PFCHA 21212113). We thank Simon Hofer and Marinus Thein for initial work on RfaH^CC^ and Dmitri Svetlov for discussions and editing.

## AUTHOR CONTRIBUTIONS

P.K.Z purified and characterized proteins and prepared complexes for CryoEM and prepared figures. N.S. prepared complexes for CryoEM. B.W. purified mutant RNAP variants and performed *in vitro* pause assays. T.H. performed cryo-EM experiments. P.K.Z. and S.H.K. carried out model building and refinement of the CryoEM structures, assisted by B.L. J. G.-H. and C.A.R.-S. performed MD simulations. All authors contributed to the data analysis. I.A. and S.H.K. wrote the first draft and revised the manuscript with contributions from C.A.R.-S., G.A.B. and M.C.W. All authors participated in preparation of the final manuscript.

## INCLUSION AND DIVERSITY

One or more of the authors of this paper self-identifies as an underrepresented ethnic minority in their field of research.

