## Supplemental Material for "Concerted transformation of a hyper-paused transcription complex and its reinforcing protein"

**Table S1.** Cryo-EM data collection, refinement, and validation statistics for complexes assembled on the *ops*PEC scaffold

| | ***ops*PEC** (PDB 8PDY) (EMDB 17626) | ***ops*PEC$^{Enc}$** (PDB 8PIB) (EMDB 17679) | ***ops*PEC$^{Rec}$ Structure 1** (PDB 8PHK) (EMDB 17668) | ***ops*PEC$^{Rec}$ Structure 2** (PDB 8PEN) (EMDB 17632) | ***ops*PEC$^{Back}$** (PDB 8PID) (EMDB 17681) | **opsPEC$^{Rec}$ + NusA** (PDB 8PIL) (EMDB 17685) |
|---|---|---|---|---|---|---|
| **Data collection and processing** | | | | | | |
| Microscope | FEI Titan Krios G3i | | | | | |
| Voltage (keV) | 300 | | | | | |
| Camera | Falcon 3EC | | | | | |
| Magnification (nominal/calibrated) | 96,000 | 96,000 | 96,000 | | 96,000 | 96,000 |
| Pixel size at detector (Å/pixel) | 0.832 | 0.832 | 0.832 | | 0.832 | 0.832 |
| Total electron exposure ($e^-/Å^2$) | 42 | 42 | 42 | | 42 | 42 |
| Exposure rate ($e^-$/pixel/s) | 0.7 | 0.7 | 0.7 | | 0.7 | 0.7 |
| No. of frames collected during exposure | 33 | | | | | |
| Defocus range (µm) | 0.80 - 2 | | | | | |
| Automation software | EPU 2.8.1 | | | | | |
| Micrographs collected (no.) | 3,790 | 5,094 | 3,382 | | 3,341 | 4,226 |
| Micrographs used (no.) | 3,613 | 4,986 | 2949 | | 2,823 | 3,142 |
| Total extracted particles (no.) | 759,854 | 1,090,135 | 1,040,421 | | 360,161 | 877,027 |
| Final particles (no.) | 250,888 | 848,008 | 124,233 | | 297,862 | 41,797 |
| Point-group or helical symmetry parameters | C1 | C1 | C1 | | C1 | C1 |
| Resolution (global, Å) | | | | | | |
| FSC 0.143 (unmasked/masked) | 4 / 3.5 | 3.2 / 2.6 | 3.7 / 3.1 | | 3.5 / 3 | 4 / 3.2 |
| Resolution range (local, Å) | 2.8 – 30.00 | 1.95 – 30.00 | 2.5 – 30.00 | | 2.2 – 30.00 | 2.4 – 30.00 |
| Map sharpening *B* factor ($Å^2$) / (*B* factor range) | -130 | -85 | -95 | | -100 | -78 |
| Map sharpening methods | local B-factor | | | | | |
| Refinement package | PHENIX (1.20_44591) real.space.refine | | | | | |
| **Model composition** | | | | | | |
| Non-hydrogen atoms | 26,887 | 27,850 | 28,225 | 27,734 | 28,297 | 30,440 |
| protein residues | 3,221 | 3,316 | 3,370 | 3309 | 3,371 | 3,656 |
| DNA nucleotides | 70 | 80 | 76 | 76 | 76 | 72 |
| RNA nucleotides | 12 | 12 | 12 | 12 | 14 | 12 |
| $Mg^{2+}$ ion | 1 | 1 | 1 | 1 | 1 | 1 |
| $Zn^{2+}$ ions | 2 | 1 | 2 | 2 | 2 | 2 |

| | | | | | | |
|---|---|---|---|---|---|---|
| **Model Refinement** | | | | | | |
| **Model-Map scores** | | | | | | |
| - CC (mask) | 0.87 | 0.88 | 0.86 | 0.87 | 0.88 | 0.84 |
| - CC (volume) | 0.86 | 0.87 | 0.85 | 0.86 | 0.87 | 0.82 |
| - Average FSC (unmasked/masked) | | | | | | |
| **Average grouped B factors (Å$^2$)** | | | | | | |
| overall | 161 | 116 | 125 | 140 | 94 | 121 |
| protein residues | 159 | 111 | 116 | 135 | 93 | 120 |
| DNA nucleotides | 198 | 200 | 280 | 179 | 113 | 140 |
| RNA nucleotides | 154 | 78 | 133 | 112 | 97 | 104 |
| $Mg^{2+}$ ion | 154 | 104 | 97 | 153 | 81 | 133 |
| $Zn^{2+}$ ions | 135 | 127 | 137 | 175 | 106 | 131 |
| **r.m.s.d. from ideal values** | | | | | | |
| Bond lengths (Å) | 0.002 | 0.003 | 0.002 | 0.002 | 0.002 | 0.002 |
| Bond angles (°) | 0.547 | 0.568 | 0.517 | 0.570 | 0.477 | 0.484 |
| **Validation** | | | | | | |
| MolProbity score | 1.68 | 2.17 | 1.59 | 1.64 | 1.49 | 1.60 |
| CaBLAM outliers (%) | 1.76 | 1.95 | 1.80 | 1.71 | 2.04 | 2.02 |
| Clashscore | 10.44 | 9.35 | 9.15 | 9.34 | 7.35 | 9.19 |
| Poor rotamers (%) | 0.22 | 2.6 | 0.1 | 1.3 | 0.3 | 0.2 |
| C-beta deviations | 0 | 0 | 0 | 0 | 0 | 0 |
| EMRinger score | 1.41 | 1.85 | 1.19 | 1.55 | 1.58 | 1.58 |
| **Ramachandran plot** | | | | | | |
| Favored (%) | 97.2 | 97.3 | 97.5 | 97.8 | 97.6 | 97.4 |
| Allowed (%) | 2.8 | 2.6 | 2.5 | 2.2 | 2.4 | 2.6 |
| Outliers (%) | 0.0 | 0.1 | 0.0 | 0.0 | 0.0 | 0.0 |
| **Ramachandran plot *Z*-score, (r.m.s.d.)** | | | | | | |
| whole | 0.68 (0.15) | 0.93 (0.15) | 1.00 (0.15) | 0.86 (0.15) | 1.18 (0.15) | 1.07 (0.14) |
| helix | 1.68 (0.16) | 1.98 (0.16) | 2.07 (0.16) | 1.77 (0.16) | 2.38 (0.16) | 2.08 (0.15) |
| sheet | 0.11 (0.25) | 0.52 (0.24) | 0.24 (0.23) | 0.45 (0.23) | 0.38 (0.24) | 0.32 (0.23) |
| loop | -0.28 (0.15) | -0.33 (0.15) | -0.16 (0.15) | -0.22 (0.15) | -0.16 (0.15) | -0.08 (0.15) |

**Table S2.** Cryo-EM data collection, refinement, and validation statistics for complexes assembled on the nc-*ops*PEC scaffold

| | **nc-*ops*PEC** (PDB 8PH9) (EMDB 17657) | **nc-*ops*PEC$^{Enc}$** (PDB 8PFG) (EMDB 17646) | **nc-*ops*PEC$^{Rec}$ Structure 1** (PDB 8PIM) (EMDB 17686) | **nc-*ops*PEC$^{Rec}$ Structure 2** (PDB 8PFJ) (EMDB 17647) |
|---|---|---|---|---|
| **Data collection and processing** | | | | |
| Microscope | FEI Titan Krios G3i | | | |
| Voltage (keV) | 300 | | | |
| Camera | Falcon 3EC | | | |
| Magnification (nominal/calibrated) | 96,000 | 96,000 | 96,000 | |
| Pixel size at detector (Å/pixel) | 0.832 | 0.832 | 0.832 | |
| Total electron exposure ($e^{-}/Å^2$) | 42 | 42 | 42 | |
| Exposure rate ($e^{-}$/pixel/s) | 0.7 | 0.7 | 0.7 | |
| No. of frames collected during exposure | 33 | | | |
| Defocus range (µm) | 0.60 - 2 | | | |
| Automation software | EPU 2.8.1 | | | |
| Micrographs collected (no.) | 4,332 | 5,230 | 4,614 | |
| Micrographs used (no.) | 4,261 | 4,979 | 4,449 | |
| Total extracted particles (no.) | 510,420 | 1,519,652 | 849,514 | |
| Final particles (no.) | 242,435 | 509,867 | 51,791 | |
| Point-group or helical symmetry parameters | C1 | C1 | C1 | |
| Resolution (global, Å) | | | | |
| FSC 0.143 (unmasked/masked) | 3.5 / 3 | 3.5 / 3.1 | 4 / 3.4 | |
| Resolution range (local, Å) | 2.54 – 30.00 | 2.48 – 30.00 | 2.8 – 30.00 | |
| Map sharpening *B* factor ($Å^2$) / (*B* factor range) | -90 | -96 | -73 | |
| Map sharpening methods | local B-factor | | | |
| Refinement package | PHENIX (1.20_44591) real.space.refine | | | |
| **Model composition** | | | | |
| Non-hydrogen atoms | 26,957 | 27,809 | 28,232 | 27,856 |
| protein residues | 3,218 | 3,311 | 3,374 | 3,313 |
| DNA nucleotides | 72 | 80 | 70 | 76 |
| RNA nucleotides | 15 | 12 | 17 | 17 |
| $Mg^{2+}$ ion | 1 | 1 | 1 | 1 |
| $Zn^{2+}$ ions | 2 | 1 | 2 | 2 |
| **Model Refinement** | | | | |
| **Model-Map scores** | | | | |
| - CC (mask) | 0.90 | 0.88 | 0.87 | 0.86 |

| | | | | |
|---|---|---|---|---|
| - CC (volume) | 0.89 | 0.87 | 0.86 | 0.86 |
| - Average FSC (unmasked/masked) | | | | |
| **Average grouped B factors (Å$^2$)** | | | | |
| overall | 116 | 121 | 151 | 151 |
| protein residues | 111 | 116 | 147 | 146 |
| DNA nucleotides | 185 | 208 | 184 | 245 |
| RNA nucleotides | 178 | 115 | 287 | 104 |
| $Mg^{2+}$ ion | 101 | 135 | 152 | 145 |
| $Zn^{2+}$ ions | 168 | 113 | 199 | 175 |
| **r.m.s.d. from ideal values** | | | | |
| Bond lengths (Å) | 0.002 | 0.003 | 0.007 | 0.002 |
| Bond angles (°) | 0.476 | 0.449 | 0.657 | 0.529 |
| **Validation** | | | | |
| MolProbity score | 1.58 | 2.21 | 1.92 | 1.93 |
| CaBLAM outliers (%) | 1.57 | 1.90 | 1.83 | 1.95 |
| Clashscore | 7.61 | 9.04 | 9.2 | 9.21 |
| Poor rotamers (%) | 1.5 | 5.5 | 2.7 | 2.94 |
| C-beta deviations | 0 | 0 | 0 | 0 |
| EMRinger score | 2.02 | 1.92 | 1.28 | 1.50 |
| **Ramachandran plot** | | | | |
| Favored (%) | 97.8 | 97.1 | 97.5 | 97.6 |
| Allowed (%) | 2.2 | 2.9 | 2.5 | 2.4 |
| Outliers (%) | 0.0 | 0.0 | 0.0 | 0.0 |
| **Ramachandran plot *Z*-score, (r.m.s.d.)** | | | | |
| whole | 1.39 (0.15) | 0.76 (0.15) | 1.21 (0.15) | 1.43 (0.15) |
| helix | 2.36 (0.16) | 1.82 (0.15) | 2.30 (0.16) | 2.28 (0.16) |
| sheet | 0.66 (0.24) | -0.01 (0.24) | 0.71 (0.24) | 0.67 (0.24) |
| loop | 0.03 (0.15) | -0.28 (0.16) | -0.20 (0.15) | 0.15 (0.16) |

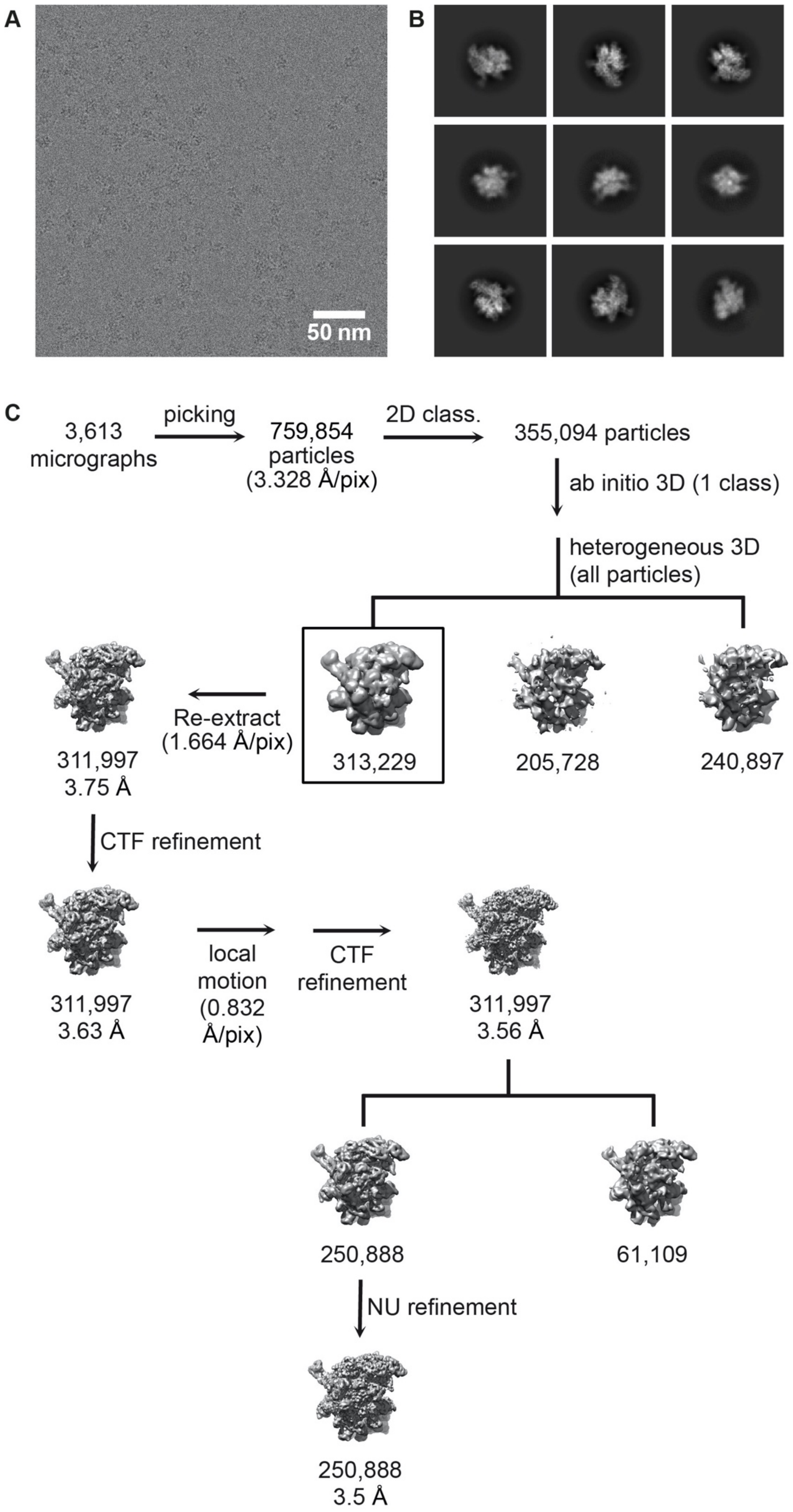

A
50 nm
B
C
3,613 micrographs
picking
759,854 particles (3.328 Å/pix)
2D class.
355,094 particles
ab initio 3D (1 class)
heterogeneous 3D (all particles)
313,229
205,728
240,897
Re-extract (1.664 Å/pix)
311,997
3.75 Å
CTF refinement
311,997
3.63 Å
local motion (0.832 Å/pix)
CTF refinement
311,997
3.56 Å
250,888
61,109
NU refinement
250,888
3.5 Å

**Figure S1. CryoEM data acquisition and processing.**

Representative micrograph (A), 2D class averages (B) and data processing flowchart (C) of the *ops*PEC cryoEM data acquisition and processing. The micrographs, 2D classes and workflow are highly similar for the samples of all presented structures and thus serve as representative example. The scale bar in (A) represents 50 nm. Class averages in (B) were generated using reference-free 2D classification within cryoSPARC. (C) A total of 3,613 aligned movies were selected for further analysis. Initially picked particles (759,854) were extracted fourier-cropped to a pixel size of 3.328 Å and subjected to reference-free 2D classification. Shiny 2D class averages were selected for ab initio 3D reconstruction to generate an initial reference for heterogeneous 3D refinement of the whole dataset. The best appearing class (boxed) was re-extracted with reduced fourier cropping and subjected to homogeneous and CTF refinement before local motion correction was conducted. Final particle images were selected by 3D heterogeneous refinement after another cycle of homogeneous and CTF refinement. Non-uniform refinement of 250,888 selected particle images resulted in a final reconstruction at a global resolution of 3.5 Å.

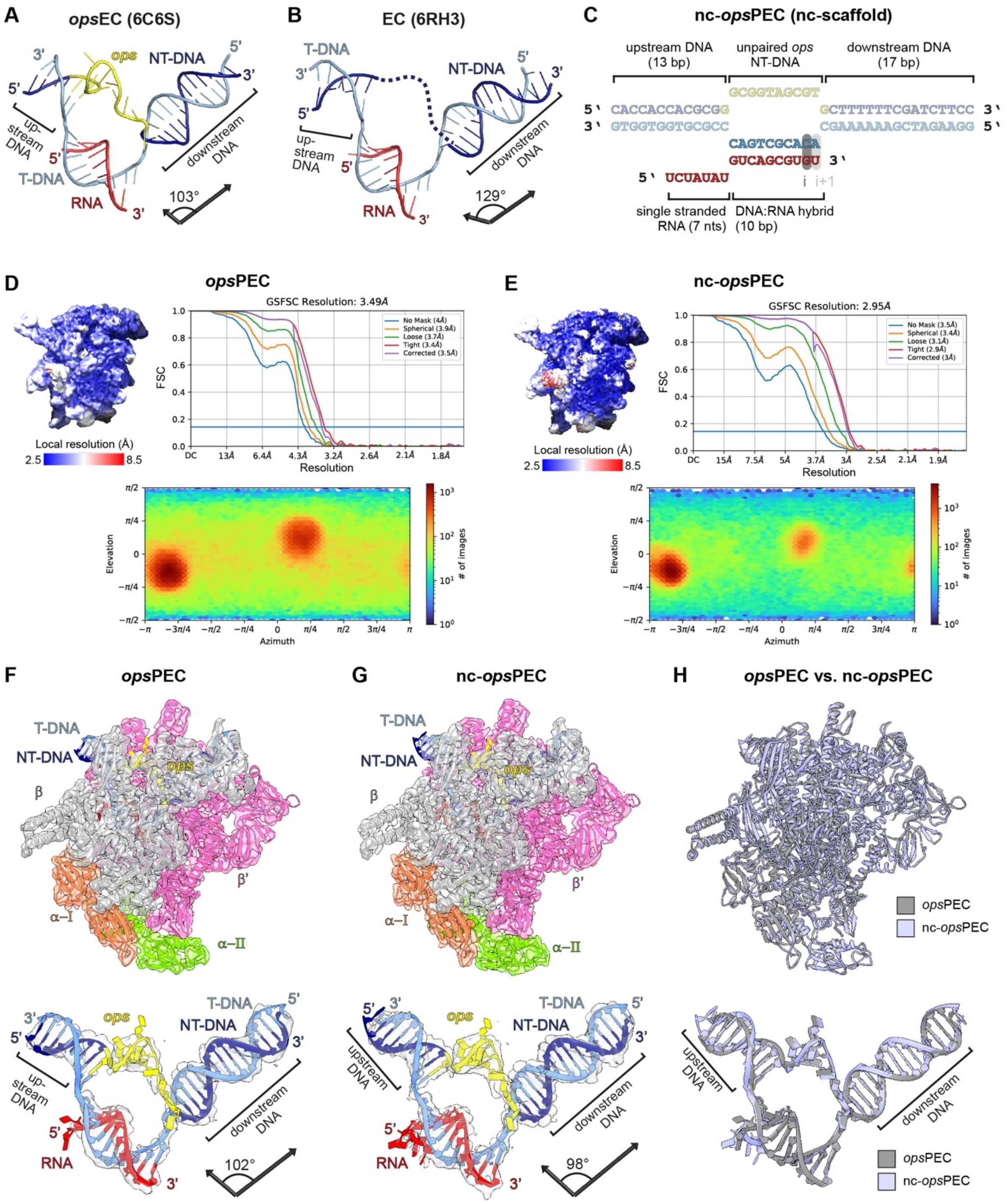


**Figure S2. Structural comparison of the *ops*PEC with other ECs and the nc-*ops*PEC.**

(A and B) Nucleic acid scaffolds of a pre-translocated EC (A, PDB-ID: 6RH3) and the *ops*EC (B, PDB-ID: 6C6S), all in cartoon representation. Helix vectors of the up- and downstream DNA (labelled) and angle between them are shown.

(C) Nucleic acid scaffold used for reconstitution of the pre-translocated nc-*ops*PEC, (harbouring a non-complementary transcription bubble). Regions identical to the scaffold of *ops*PEC (compare Figure 1B) are greyed out, divergent regions are shown in solid colour.

(D and E) CryoEM data statistics for *ops*PEC (D) and nc-*ops*PEC (E). The local resolutions (top left), Fourier shell correlation (FSC, top right) and particle angular distributions plots after NU refinement (bottom) are displayed. Local resolutions range from 2.5 Å (blue) to 8.5 Å (red); low resolution regions mainly reside within the upstream DNA and flexible RNAP domains (e.g. SI2 or SI3).

(F – H) Structural comparison of *ops*PEC and nc-*ops*PEC. Top: Models of *ops*PEC (F), nc-*ops*PEC (G) and their superposition (H) are shown as cartoon along with their cryoEM maps (F and G; transparent surface). RNAP and corresponding maps are coloured according to RNAP subunits, nucleic acid density is light blue (F, G). Bottom: Nucleic acid scaffolds of *ops*PEC (F) or nc-*ops*PEC (G) together with their cryoEM densities (transparent surface) and their superposition (H). In (F) and (G), the helix axis vectors of up- and downstream DNA and the angle between them are shown. In (H), structures are coloured as indicated.

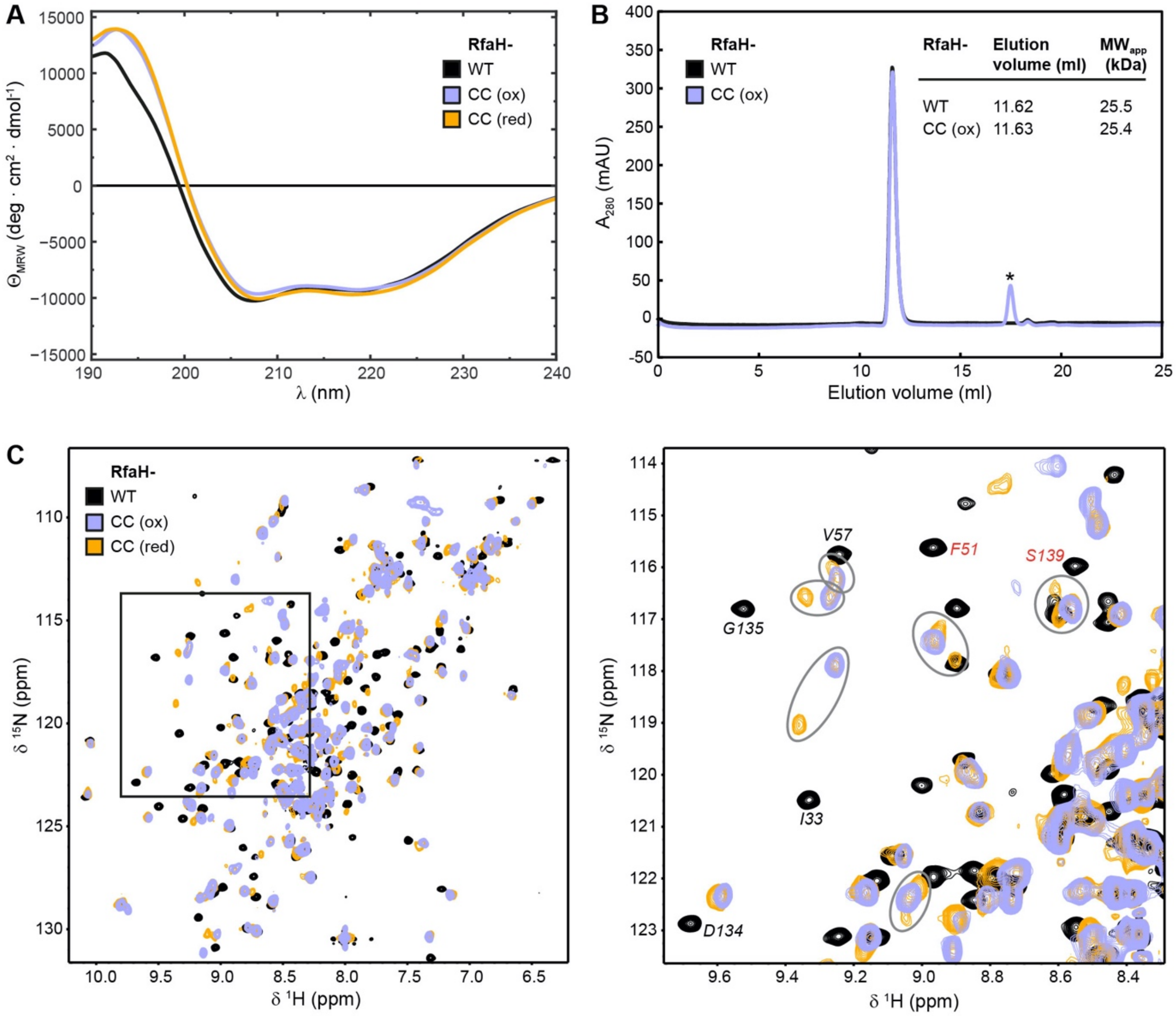


**Figure S3. Protein quality control of RfaH$^{CC}$.**

(A) Overlay of normalized CD-spectra of WT-RfaH (black) and oxidized RfaH$^{CC}$ (purple), recorded in CD buffer (10 mM $KP_i$ (pH 7.0)) and of RfaH$^{CC}$ in CD buffer containing 0.5 mM TCEP (orange).

(B) Overlay of chromatograms of analytical SEC runs of WT-RfaH (black) and oxidized RfaH$^{CC}$ (purple) on a 24 ml Superdex 75 column. The asterisk marks a buffer artefact. The inset shows the elution volumes determined for the main peaks and the proteins' apparent molecular weights calculated by comparison of the elution volumes of standard proteins (see STAR Methods).

(C) Left: overlay of 2D [$^{1}H$, $^{15}N$]-HSQC spectra of $^{15}N$-labelled WT-RfaH (black) and $RfaH^{CC}$ in the presence of $Cu^{II}$ phenanthroline (oxidized state; purple) or DTT (reduced state; orange), respectively. Right: enlargement of the boxed spectral window of the full spectrum (left). Resonances of the original RfaH-F51 and RfaH-S139 backbone amides disappearing due to the substitutions and of those in spatial proximity to the substitution sites are labelled in red or black, respectively. Ellipses highlight signals that are split in two or more (weak) peaks in the reduced state (indicative of multiple different conformations) but reduce to one (strong) signal in the oxidized state (indicative of fixing one particular conformation by cystine formation).

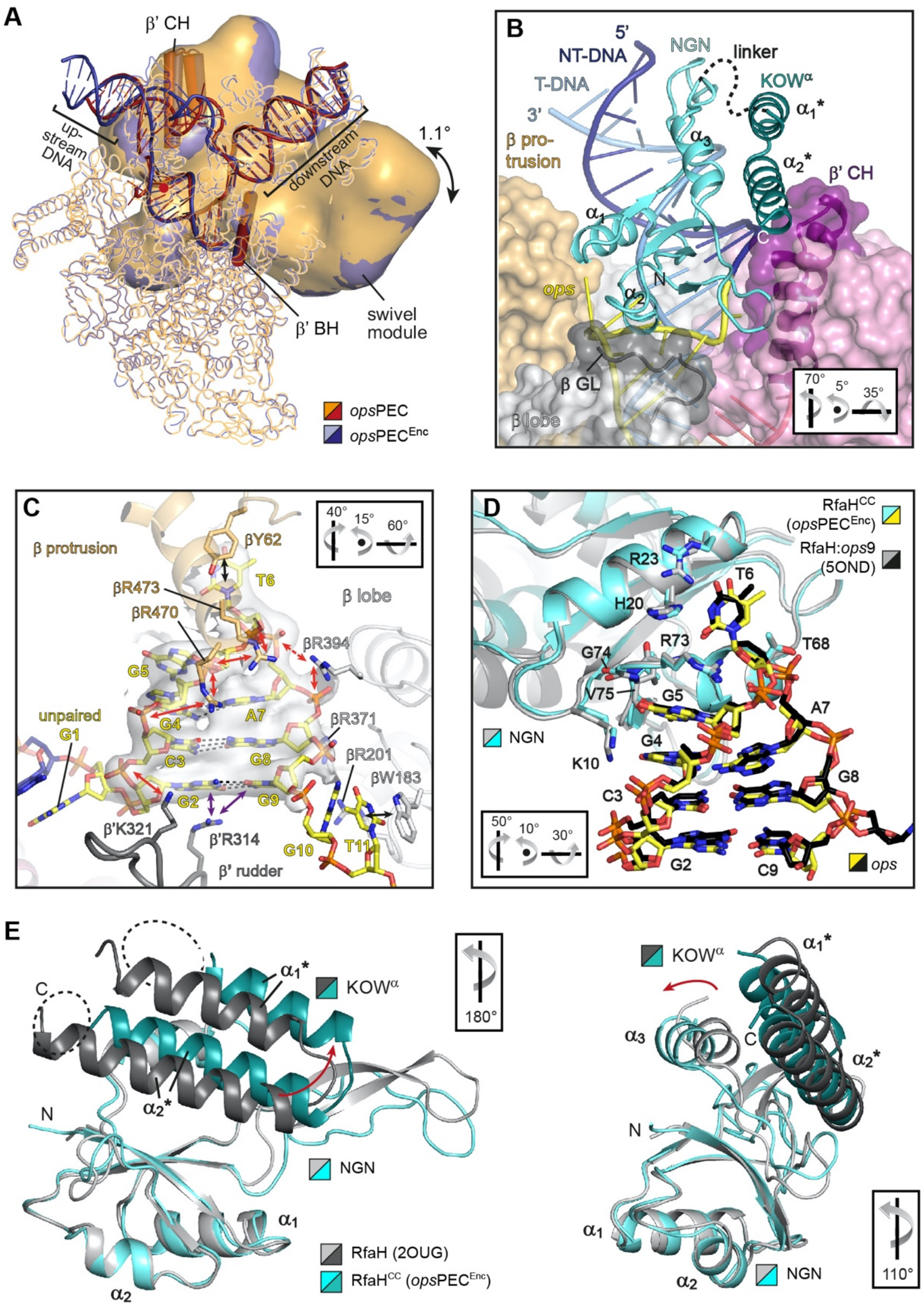


**Figure S4. Structural analysis and comparison of the *ops*PEC$^{Enc}$ complex.**

(A) *ops*PEC$^{Enc}$ is less swivelled than *ops*PEC. Shown is a structural overlay of the *ops*PEC and *ops*PEC$^{Enc}$. The two models are superimposed on the core region (ribbons) with the swivel module being represented as Gaussian surface. Swivel angle and approximate axis (red dot) are indicated. The $\beta$’ BH and $\beta$’ CH are shown as cartoon tubes to indicate the orientation of the swivelling axis.

(B) The RfaH$^{CC}$ KOW$^{a}$ domain is located on top of the $\beta$’ CH. RNAP is depicted as molecular surface; selected structural elements are shown as cartoons and coloured as indicated. RfaH$^{CC}$ (cyan/mint) is in cartoon representation, relevant secondary structure elements are labelled. The orientations relative to the standard view (Figure S4A) are indicated.

(C) Accommodation of the *ops*HP within the *ops*PEC$^{Enc}$ complex. *ops* DNA is in stick representation, selected RNAP elements are in cartoon with side chains of residues contacting *ops* shown as sticks. H-bonds of the *ops*HP base pairs are represented by dashed lines, stacking interaction by arrows. The CryoEM map of the *ops*HP is shown as transparent surface. The orientation relative to the standard view (Figure S4A) is indicated.

(D) Recognition of *ops* by RfaH in *ops*PEC$^{Enc}$ is the same as in bimolecular RfaH:*ops* complex. Superposition of the RfaH:*ops*9 complex of a co-crystal structure (PDB-ID: 5OND) and RfaH$^{CC}$:*ops*HP of the *ops*PEC$^{Enc}$ complex. RfaH in cartoon representation, side chains of *ops* interacting residues and *ops* DNAs shown as sticks. Colors are as indicated. The orientation relative to the standard view (Figure S4A) is given.

(E) The RfaH$^{CC}$ KOW$^{\alpha}$ helices shift position upon binding to the *ops*PEC and unfold partially. Overlay of free RfaH (PDB-ID: 2OUG, cyan/mint) and RfaH$^{CC}$ of the *ops*PEC$^{Enc}$ (light/dark grey). Alpha helices and termini are labelled. (Left) Shift of the KOW$^{\alpha}$ helices away from the NGN, and the unfolding of the C-terminal $\alpha_1$* and $\alpha_2$* helix turns. (Right) NGN helix $\alpha_3$ is pushed together with the KOW$^{\alpha}$. The orientations relative to the standard view (Figure S4A) are indicated.

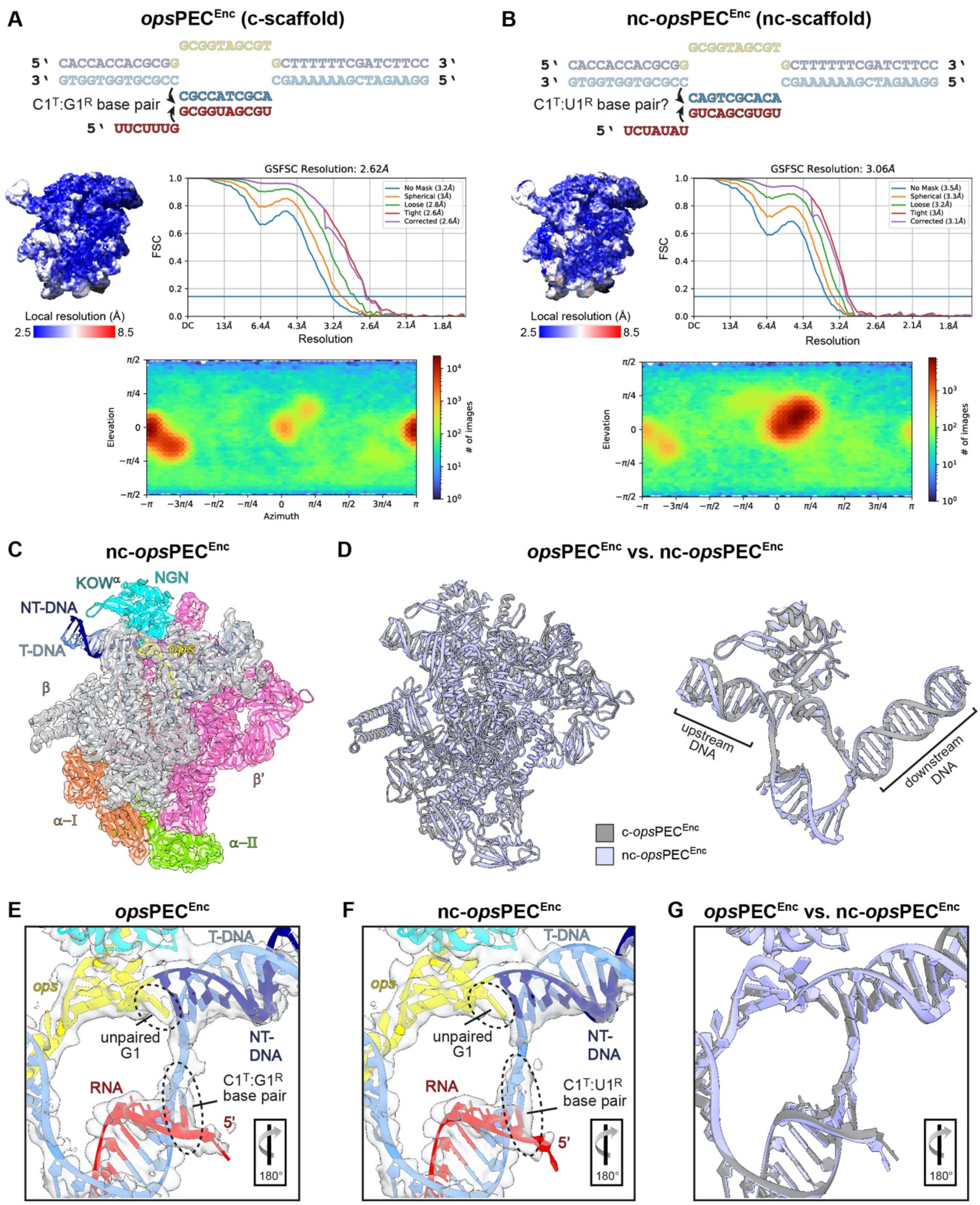


**Figure S5. Comparison of the *ops*PEC$^{Enc}$ and nc-*ops*PEC$^{Enc}$ structures.**

(A and B) Top: Nucleic acids scaffolds used for reconstitution of the *ops*PEC$^{Enc}$ (A) or nc-*ops*PEC$^{Enc}$ complexes (B). Regions identical in both scaffolds are greyed out, divergent regions are shown in solid colour. The arrows show formation of the C1$^{T}$:G1$^{R}$ base pair in *ops*PEC upon binding of RfaH$^{CC}$ (A), or the formation of a non-canonical C1$^{T}$:U1$^{R}$ base pair in the non-complementary scaffold. Middle left: Local resolutions of the respective complexes range from 2.5 Å (blue) to 8.5 Å (red); low resolution regions mainly reside within the upstream DNA and flexible RNAP domains (e.g. SI2 or SI3). Middle right: Fourier shell correlation plots of the two complexes. Bottom: particle angular distribution plots.

(C) CryoEM map (transparent surface) and model (in cartoon representation) of the nc-*ops*PEC$^{Enc}$ complex. Colours as indicated.

(D) Overlay of *ops*PEC$^{Enc}$ and nc-*ops*PEC$^{Enc}$ structures. Left: Whole models, right: nucleic acids and RfaH$^{CC}$ only. Models are shown as cartoon with colours as indicated.

(E - G) Close-up view of the upstream edge of the transcription bubble. Nucleic acid scaffods (cartoon representation) and their associated cryoEM density (transparent surface) are shown for *ops*PEC$^{Enc}$ (E), nc-*ops*PEC$^{Enc}$ (F) and an overlay of the two structures (G). The G1 base is unpaired in both structures and a 11$^{th}$ RNA:T-DNA hybrid base pair at its upstream end is formed instead (C1$^{T}$:G1$^{R}$ in *ops*PEC$^{Enc}$ and a non-canonical C1$^{T}$:U1$^{R}$ in nc-*ops*PEC$^{Enc}$). The orientation relative to Figure S5C is indicated.

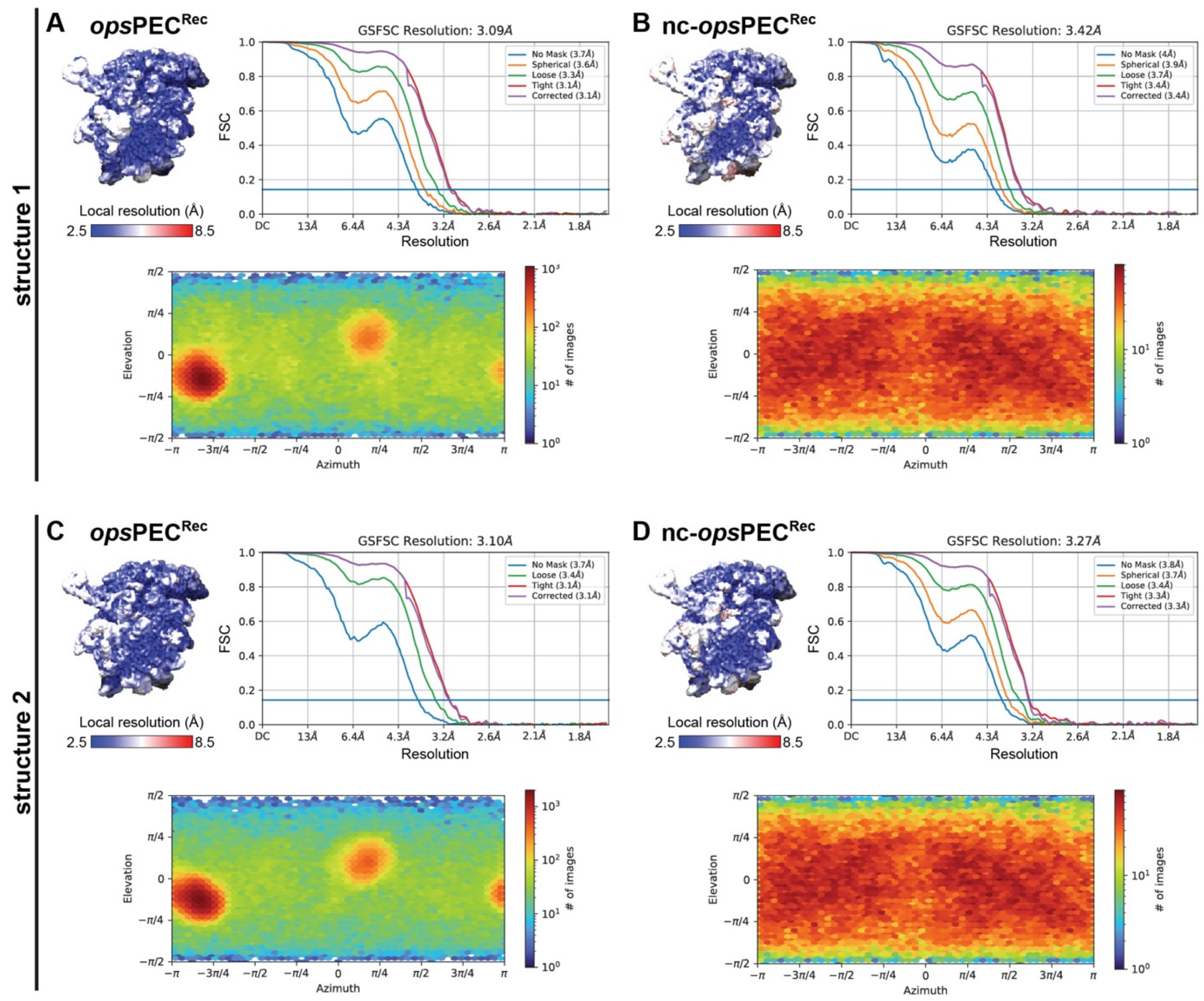


**Figure S6. Analysis of the cryoEM data sets of *ops*PEC$^{Rec}$ and nc-*ops*PEC$^{Rec}$ structures 1 and 2.**

(A – D) Top left: Local resolutions of the complexes plotted on their corresponding cryoEM densities. The local resolutions range from 2.5 Å (blue) to 8.5 Å (red). Low resolution regions mainly reside within the upstream DNA and flexible RNAP domains. Top right, Fourier shell correlation plots and, bottom, particle angular distribution plots of the corresponding complexes. Structures 1 and 2 represent the two states obtained from 3DVA of the *ops*PEC$^{Rec}$ or nc-*ops*PEC$^{Rec}$ complexes, respectively.

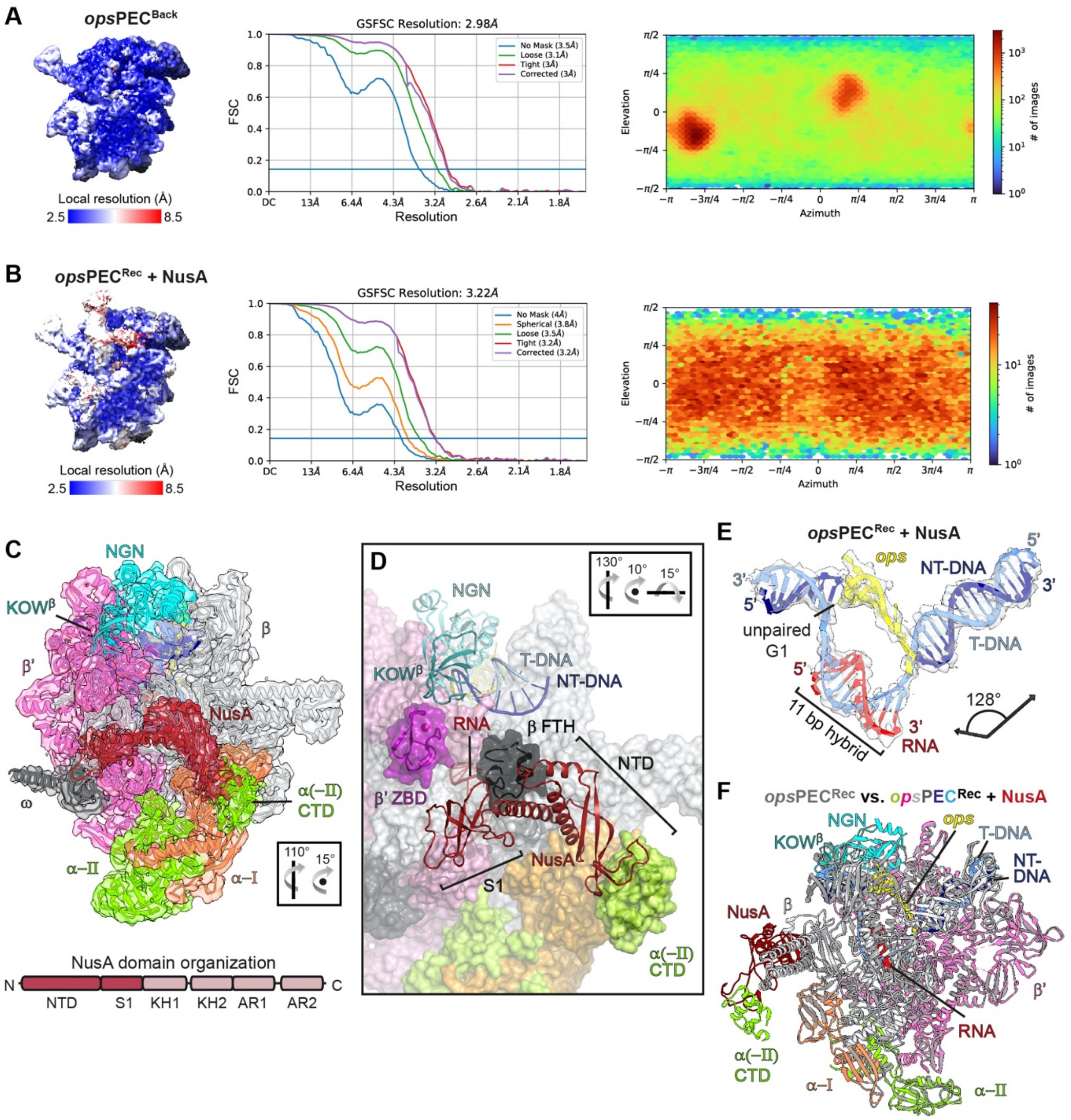


**Figure S7. CryoEM data and map quality parameters of the *ops*PEC$^{Back}$ and *ops*PEC$^{Rec}$:NusA complexes, and structural details of the NusA modified complex.**

(A and B) Left: Local resolutions of the *ops*PEC$^{Back}$ (A) or *ops*PEC$^{Rec}$ + NusA (B) complexes plotted on their corresponding cryoEM densities. Local resolutions range from 2.5 Å (blue) to 8.5 Å (red). Low resolution regions mainly reside within the upstream DNA and flexible RNAP domains. Middle, Fourier shell correlation plots and, right, particle angular distribution plots of the corresponding complexes.

(C) Overview of the *ops*PEC$^{Rec}$ + NusA complex. The cryoEM map of the complex is shown along with the model, both colour-coded. The domain organization of NusA is depicted at the bottom; greyed-out domains are not resolved in the map, i.e. only NusA-NTD and S1 domains are visible. The orientation relative to the standard view (Figure 2A) is indicated.

(D) Close-up view of the NusA:RNAP interactions. RNAP is in surface representation, domains interacting with either NusA ($\beta$ FTH and $\alpha$-CTD) or RfaH KOW$^{\beta}$ domain ($\beta$' ZBD) are shown as cartoon (the $Zn^{2+}$ ion of the $\beta$' ZBD is shown as sphere). NusA is displayed in cartoon representation, domains are labelled. The orientation relative to the standard view (Figure 2A) is indicated.

(E) Structure of the *ops*PEC$^{Rec}$ + NusA nucleic acid scaffold with associated cryoEM densities. Nucleic acid strands are shown as cartoon, the DNA's/RNA's orientations are indicated. The arrows represent the helix vector of the up- and downstream duplexes, the angle between within them is indicated.

(F) NusA binding to the *ops*PEC$^{Rec}$ does not alter the conformation of the RNAP, nucleic acids or RfaH. Superposition of the *ops*PEC$^{Rec}$ (grey) and the *ops*PEC$^{Rec}$ + NusA complexes (coloured as indicated), both depicted in cartoon representation.
